# Intramolecular Communication and Allosteric Sites in Enzymes Unraveled by Time-Dependent Linear Response Theory

**DOI:** 10.1101/677617

**Authors:** Bang-Chieh Huang, Yi-Yun Cheng, Chi-Hong Chang-Chein, Kuan-Chou Chen, Chao-Ling Yao, Lee-Wei Yang

## Abstract

It has been an established idea in recent years that protein is a physiochemically connected network. Allostery, understood in this new context, is a manifestation of residue communicating between remote sites in this network, and hence a rising interest to identify functionally relevant communication pathways and the frequent communicators within. However, there have been limited computationally trackable general methods to discover proteins’ allosteric sites in atomistic resolution with good accuracy. In this study, we devised a time-dependent linear response theory (td-LRT) integrating intrinsic protein dynamics and perturbation forces that excite protein’s temporary reconfiguration at the non-equilibrium state, to describe atom-specific time responses as the propagating mechanical signals and discover that the most frequent remote communicators can be important allosteric sites, mutation of which could deteriorate the hydride transfer rate in DHFR by 3 orders. The preferred directionality of the signal propagation can be inferred from the asymmetric connection matrix (CM), where the coupling strength between a pair of residues is suggested by their communication score (CS) in the CM, which is found consistent with experimentally characterized nonadditivity of double mutants. Also, the intramolecular communication centers (ICCs), having high CSs, are found evolutionarily conserved, suggesting their biological importance. We also identify spatially clustered top ICCs as the newly found allosteric site in ATG4B. Among 2016 FDA-approved drugs screened to target the site, two interacting with the site most favorably, confirmed by MD simulations, are found to inhibit ATG4B biochemically and be tumor suppressive in colorectal, pancreatic and breast cancer cell lines with an observed additive therapeutic effect when co-used with an active-site inhibitor.

## INTRODUCTION

The protein structures, dynamics and functions have been intensively studied for several decades. One of the long-standing challenges in revealing the structures-dynamics-function relationship is to estimate the synergistic effects of remote sites, such as relatively buried enzyme active site and alternative binding sites for activators/inhibitors – known as the allosteric effect (1-3). Since 60s, pre-existing model (MWC model) (4) and induced fit model (KNF model) (5) have been used to describe such a mechanism driven by noticeable conformational changes upon ligand binding at a non-orthosteric site. However, it has been first proposed in 1984 by Cooper and Dryden (64) and recently confirmed by NMR experiments (6) that remote mutations could impact the activity of functional sites without involving notable conformational changes. One such example is the cyclic adenosine monophosphate (cAMP) binding to the dimeric catabolite activator protein (CAP), a transcriptional activator. Kalodimos’s group used the CPMG technique (6) to quantify microsecond-millisecond dynamics in three different liganded states – no ligand, one monomer binding with a ligand, and two bound ligands in the CAP dimer while one in each monomer. It was found that the binding of one ligand actually generates microsecond-millisecond dynamics that is unseen in unliganded and fully liganded states, causing the negative cooperativity in the binding of the second cAMP binding (6).

Another example is the *E*.*coli* dihydrofolate reductase (DHFR), which catalyzes the reductation of dihydrofolate (DHF), facilitated by the cofactor NADPH, to terahydrofolate (THF) and NADP^+^ (Fig 1). The catalytic cycles (7, 8), kinetic data (9-13), folding stability (14-16), intermediate structures (17, 18), and binding dynamics (19, 20) and its hydride transfer rate of the mutants (7, 10, 11, 21-32) have been extensively studied. Recently, more evidences establish the dynamical view point of the enzyme catalytic process in the hydride transfer of DHFR (28, 30, 33, 34). One mutant G121V, 16Å away from the catalytic center, causes no noticeable structural changes (8, 20, 33) and can reduce the hydride transfer rate by 200 folds (10, 11, 26) while other remote mutants G67V (11), S148A (7, 23) and W133F (28) do not impair the activity. The QM/MM study also suggests that when a reaction progresses from the reactants to products through the transition states of the protein and/or the chemical compound, a network of “coupled promoting motions*”* could facilitate the chemistry in DHFR (12, 34). Furthermore, a recent NMR CPMG experiment showed that dynamics on the microsecond-millisecond time scales are significantly changed in the G121V mutant of DHFR, owing to the mutant inducing impaired residue network inside DHFR, which decreases dynamical signal propagation (33).

**Figure 1.**
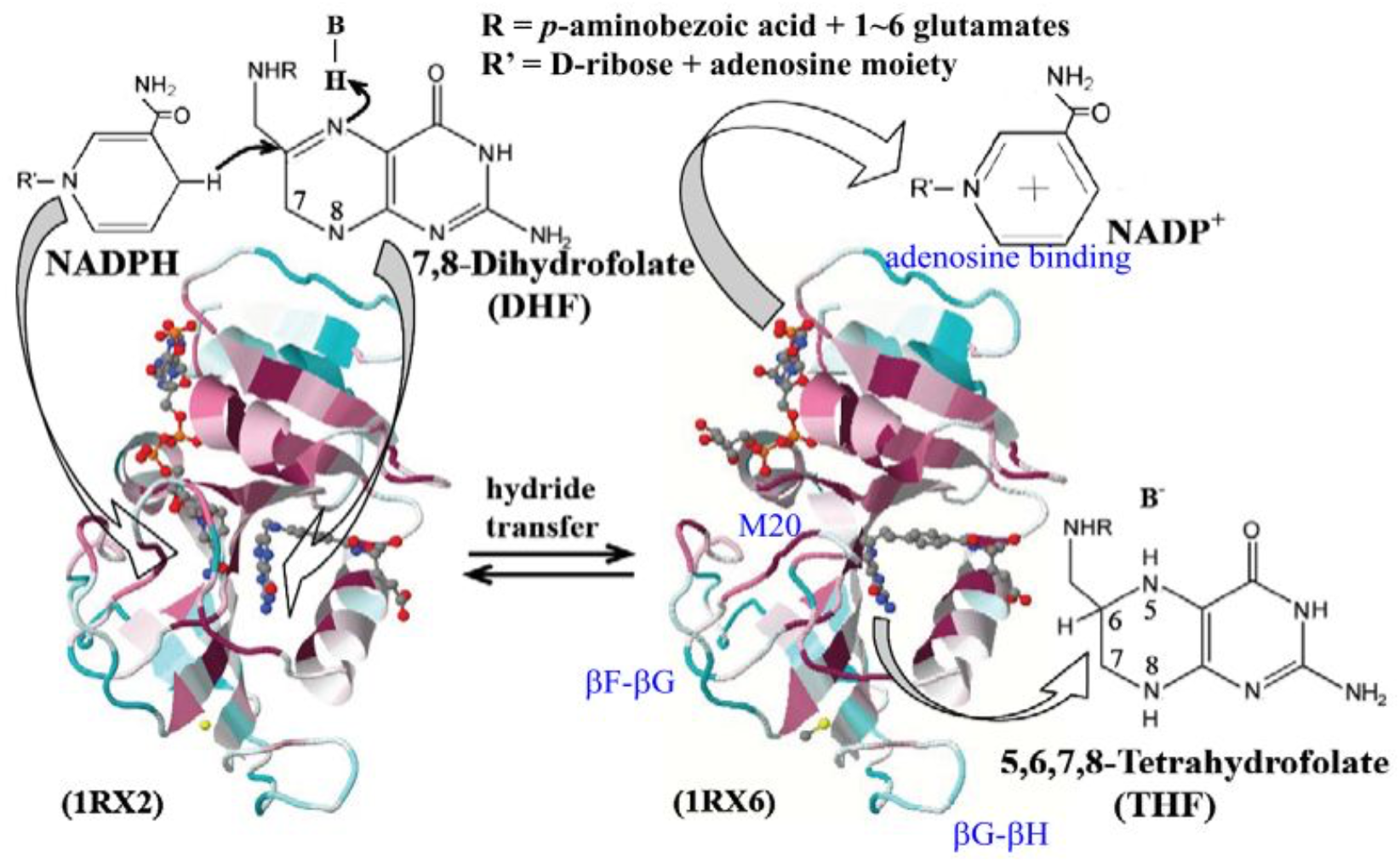
E. coli DHFR (“DHFR*”* thereafter) facilitates the hydride transfer from NADPH to the C6 of the pterin nucleus with a concurrent protonation in N5 position of DHF in the reaction of NADPH + H^+^ + DHF → NADP^+^ + THF. The hydride transfer-prompting Michaelis (binary) complex on the left (PDB ID: 1RX2) and the product ternary complex on the right (PDB ID:1RX6) are color-coded by residue’s evolutionary conservation based on multiple sequence alignment of 77 DHFR homologs from cyan, the most variable, to dark red, the most conserved using Consurf server (66-68).

Besides NMR, other spectroscopy data also suggested that low frequency collective vibrational excitations exist in *ps-ns* range could mediate certain degree of allostery (35-38). Yet, it is still an open question as for how proteins, say enzymes, exploit the vibrational excitations through the signal propagation to regulate catalysis of chemical reactions. In the past decade or two, growing number of theoretical methods were proposed to elucidate intramolecular signal propagation pathways (39-48). In general, the methods can be classified into two categories – structural/dynamical equilibrium approaches (42-44, 46, 47) and those of non-equilibrium (39-41, 45, 48). On the equilibrium approaches, Chennubhotla and Bahar used the Markov approach to extract the communicating residues in a GroEL-GroES system (42, 47). Vishveshwara’s group applied Floyd-Warshal algorithm to determine shortest paths between residues having high correlation in positional deviations, analyzed from MD trajectories (43). Kong and Karplus used an unsupervised clustering technique to group residues that are highly correlated in a PDZ domain, and the distributions of communicating residues in a given cluster could report signal ‘pathway’ (44). Despite the good efforts, one major problem of these equilibrium approaches is that the propagation pathways are fixed and the perturbations/external forces play no roles in the resolved pathways.

Non-equilibrium approaches served as promising alternatives to investigate signal propagations through the network. Ota and Agard have developed a protocol called “anisotropic thermal diffusion (ATD)*”* to study the kinetic energy propagation using a non-equilibrium molecular dynamics (MD) simulation (41). The ATD starts with cooling down the protein to 10K in vacuum then a specific residue is heat up to 300K. Consequently, the signal propagation pathway from the heated spot can be tracked by the R.M.S.D. changes of atoms deviated from their equilibrated positions as a function of time. It was found that the speed of intramolecular signaling is about 14 Å*/*ps, which is comparable to the speed of sound in liquids water at room temperature. However, ATD suffers from low signal-to-noise ratio, which makes it hard to trace the complete network through a protein; also the non-native environment is a concern. Sharp and Skinner used “pump-probe molecular dynamics simulations (PPMD)*”* to study the signal propagation. Selected residues are ‘pumped’ by oscillatory forces in a period ∼10 ps, the ‘probes’, are found within those that have the highest enrichment in Fourier-transform-derived density of states for the ‘pumped’ frequency. The phase delay divided by inter-residue distance is interpreted as the speed of the signal, found at 5Å/ps (45). Although the “mutual correlated residues” were identified by PPMD, the time order of signal propagation is absent. Leitner’s group combines the non-equilibrium MD and the normal mode relaxation process to study the diffusivity of heat current in the hemoglobin (39). Given an initial position and velocity, the normal mode relaxation method provides the displacement and velocity as a function of time, which serves as the route to estimate the kinetic energy propagation. Notably, Ranganathan group proposed a correlated mutation method called the “statistical coupling analysis (SCA)*”* to identity the co-evolved residues (46). This approach, requiring many homologous sequences to secure its statistical significance, cannot easily distinguish the structural or dynamics relevance with the found co-evolved mutation sites. An example is that the statistical coupling site W133, 20.1Å away from the hydride transfer center, shows no obvious effect on the fold reduction of hydride transfer rate in DHFR while SCA surmises its regulatory role in catalysis (28, 46). In fact, the good method, after refinement by maximum-entropy modeling, is later used for protein structure prediction from primary sequences (65). Recently, Thirumalai’s group summarized a method to predict the “allostery wiring diagram*”* (AWD) in a protein which described residue network regulate the allosteric functions (49). The AWD can be derived from the aforementioned SCA (50), structural perturbation method (SPM) using ENM or NMA (50-52) and hamiltonian switch method (HSM) (53).

Supplementing the aforementioned theories, earlier we developed a general time-dependent linear response theory (td-LRT) to investigate the time responses of protein reconfiguration subject to external perturbations (e.g. ligand dissociation) (54)(54b). The method, combining normal mode analysis (NMA) (55, 56) and damped harmonic oscillators damped in water, solved by Langevin equation (57), has been shown to reproduce the relaxation time of specific sites in carbonmonoxy myoglobin (Mb) measured by time-resolved UV Resonance Raman (UVRR) spectroscopy (36) and time-resolved X-ray crystallography (36, 58). Using td-LRT, we are able to monitor a two-staged relaxation where the slower relaxation ranges from 4.4 to 81.2 ps, while the faster ‘early responses’, ranging from hundreds of femtoseconds to a few picoseconds, can be best described by the theory when impulse forces are used. Furthermore, we identify several residues in Mb as ‘disseminators’ that propagate the signals the fastest and are found kinetically important to modulate gas molecule diffusion and rebinding (54, 59). We further did several point mutations and found that their kinetic importance can be reflected by the number of retained disseminators identified from the wild type (54). The quantitative agreement with several experimental data (36, 58, 59) sets an important distinction of our method.

In this article, we apply this td-LRT theory with impulse forces to investigate the dynamic driven allosteric regulation. We propose a new method to track the atomic-level dynamic signal propagation pathways, and consequently the frequent residue communicators as the intra-molecular communication centers (ICCs). The evolutional conservation and functional importance of these ICCs are carefully examined. The coupling between mechanical perturbations induced by chemical modifications on allosteric sites (or sites remote to the active sites) and fold of the reduction of experimentally characterized hydride transfer rates (at the active site) of DHFR is reported. Using spatially clustered ICCs with high communication scores as the newly found allosteric site in ATG4B, we screened 2016 FDA-approved drugs by small-molecule docking and found the ones most favorably interacting with the site, confirmed by MD simulations. The top drugs showed ATG4B inhibition biochemically and tumor suppression in three cancer cell lines with an observed additive therapeutic effect when co-used with an active-site inhibitor against ATG4B.

## THEORY, ALGORITHM and METHODS

### 2a. The System preparation

To study the dynamics relating the forward hydride transfer, the Michaelis-complex of DHFR (PDB code: 1RX2) including cofactor NADPH and substrate DHF is used to perform the normal mode analysis. The DHFR structure is formed by the rigid frame consist of an eight-strand β-sheet and four *α*-helices as well as three flexible loops: Met 20 loop (residues 9-24), F-G loop (residues 116-132) and G-H loop (residues 142-150) (20).The scalable molecular dynamics program, NAMD (Nanoscale Molecular Dynamics), (60) with CHARMM 36 force-field (61) is used to perform the energy minimization of full atom. Besides, the force-field of substrate, folate, is generated from the ParamChem web site (62, 63) based on the CHARMM General Force Field (CGenFF), where a further Q.M. calculation with MP2/6-31g(d) calculation for the folate support that the charges are close to the estimation of the ParamChem. After several million MD steps of energy minimization, the system reaches our criteria such that the mean force of atoms equals to 10^-5^ kcal/mol/ Å. Consequently, the full atom (*N=2615* atoms including all the hydrogen atoms) normal mode analysis (NMA) is carried out by diagonalizing the hessian matrix constructed by the second derivatives of the potential energy with respect to 3*N* mass-weighting Cartesian coordinates. Further detail steps of the NMA can be found in our previous work (54) and its supporting materials.

### 2b. The time-dependent linear response (td-LRT) theory and the characteristic time

To study a biomolecule perturbed by the impulse forces, we have developed a time dependent linear response theory (*td*-LRT) to describe the relaxation dynamics (54), which addresses dynamic allostery (64). The time progression of the positional changes of atom *i* are given by the following form

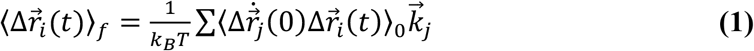

where 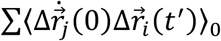is the velocity-position time-correlation function sampled in the absence of perturbations, 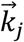are the impulse forces applying on atom *j*. Consider a protein is surrounded by a viscous environment, we express 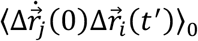as superimposed independent harmonic oscillators (55, 56, 64) under solvent damping by solving the Langevin equation (57). The derivation can be found in supporting materials of our previous work (54). Through our previous studies (54), the speed of signal propagation is a function of locations/directions of perturbation forces and the network architecture within a protein.

When impulse forces are introduced on atom *j*, atom *i* responds to these perturbations by its temporary positional departure from its equilibrium position as a function of time. In this process, the atom reaches its maximal deviation at a characteristic time 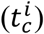 before the deviation eventually vanishes at long time (**Fig 2a**). We found that 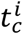is independent of the magnitude of 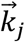but varies with the directions and location of it. To examine the robustness of the perturbation-response relationship, we define a force ensemble, *k*(Ω_*j*_) = *k*(*θ*_*j*_, *ϕ*_*j*_), acting at atom *j* with the same magnitude pointing toward 133 directions where *θ*_*j*_ is the angle between the direction of force 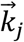and the z-axis spanning from 0° to 90° with a step size 15°, and *ϕ*_*j*_ is the angle between the projection vector of the force 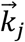on the x-y plane and the x-axis spanning from 0° to 345° with the same step size 15°. Therefore, the characteristic times corresponding to a force ensemble *k*(Ω_*j*_) are denoted by 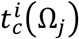.

**Figure 2.**
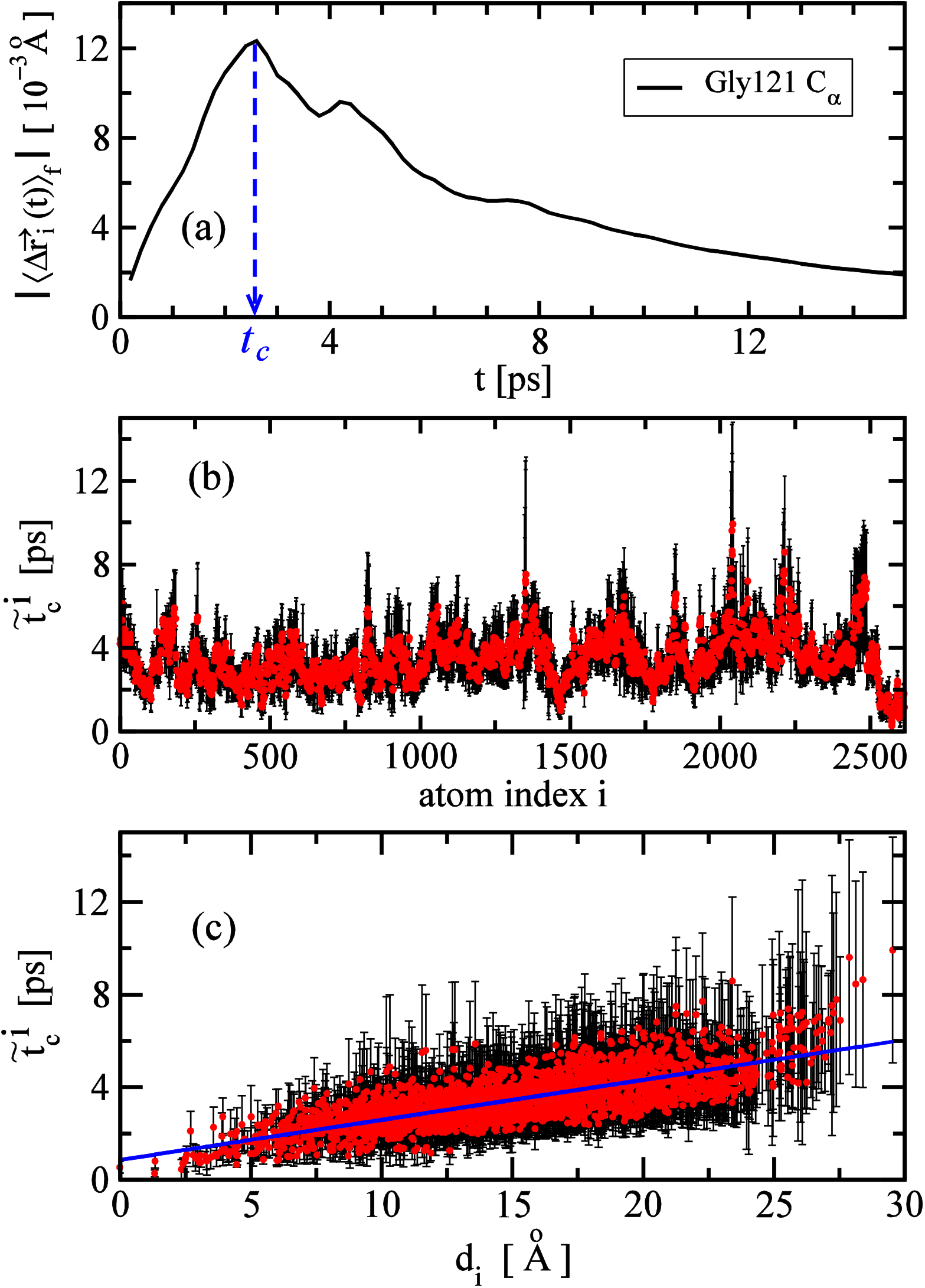
**(a)** The displacement of Gly 121 *C*_*α*_ atom is a function of the response time with the characteristic time *t*_*c*_ = 2.6 ps predicted by LRT using an impulse force on the N5 atom of the DHF. **(b)** By applying 133 evenly distributed forces on the N5 atom of the DHF, a red dot is the average characteristic times 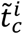 of atom *i* and a black error bar is the standard deviations. **(c)** The 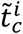as a function of the distance *d*_*i*_ between atom *i* and the N5 atom of the DHF, where the blue line is the linear regression with the correlation coefficient of 0.76. The inverse slope provided to estimate the propagation speed of 580 *m*/*s*, and the estimated speed is compatible to previous results (41, 45).

### 2c. The longest dissemination time (LDT) relating to the active sites

To understand the role of residues in mediating signals, we would like to measure how fast the signal can propagate throughout the entire enzyme from a residue; in other words, to find the longest response time for all possible signals starting from a specific residue. As in the method proposed in our previous work (54), the force ensemble defined in section 2b was applied on the *C*_*α*_ atom of residue *s*. Consequently, a set of characteristic times consist of 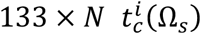can be calculated (*N* is the number of *C*_*α*_*s* or residues, which is 159 for DHFR). The longest 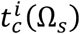in the *i*-th residue among all the 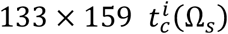values is the longest dissemination time (LDT) for residue *s*. The residues with short LDT are called the “disseminators*”* (54), suggesting their roles in efficiently broadcasting the signals throughout the protein matrix. We calculate the LDT for each residue in DHFR by perturbing the corresponding *C*_*α*_ atom, also the C6 atom of the NADPH and N5 atom of the DHF. We then compare these calculated disseminators with the active site residues reported in reference (65).

### 2d. Coarse-Grained Connection Matrix (CGCM) and communication score (CS)

Although td-LRT provides the characteristic time 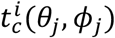to characterize the time-evolution of signal propagation, we still need a method to trace frequent signal propagation pathways and identify crucial residues along these pathways, especially when the perturbation is introduced in a number of sites along 133 directions. Here, we introduce “the Coarse-Grained Connection Matrix (CGCM)*”* to record the signal propagations between residues. The idea is illustrated in **Figure S3**. With the CGCM, we can then extract “the intramolecular communication centers (ICCs)*”* - the residues having high CSs (see below) on popular pathways communicating dynamic signals.

Because the characteristic time 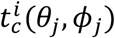provides us the causality of signal propagation, we can trace the signal transduction pathway consist of the donor-acceptor pairs. There are **two criteria** to define whether a donor atom propagates a signal to an acceptor atom, or say, whether two atoms are viewed as “connected*”* in the context of signal propagation. The atoms in the donor and acceptor residues are denoted as *atom*_*d*_ and *atom*_*a*_, respectively. **First**, given an impulse force exerted on the *j*-th *C*_*α*_ atom, 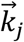, the atoms with characteristic times that differ by a small time interval Δ*t* can be viewed as “connected*”* such that 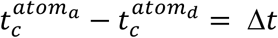, where the chosen Δ*t* is the constant time interval we store/record the time correlation functions, or equivalently, the evolving protein conformations responding to the perturbation. Δ*t* is 0.2 ps in our study. **Figure S3** demonstrates how the connection signal is recorded into a connection matrix. In order to track the physical causality of signal propagation, the “connected*”* pairs should also meet the **second** criteria - the angle between the vector *j-*th *C*_*α*_ to *atom*_*a*_ and *j-*th *C*_*α*_ to *atom*_*d*_ should be less than 90 degree. It is possible that a donor atom connects to several acceptor atoms, or several donors connect to an acceptor atom. As a result, in order to quantitatively recognize how frequently a residue participate in the signal propagation, we define a full-atom connected matrix *F*(*atom*_*d*_, *atom*_*a*_) account for the number of the connection that the donor, *a*_*d*_, connects to the acceptor, *a*_*r*_

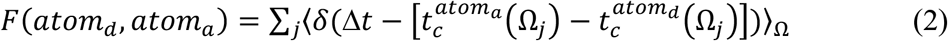

where *j* runs over all the selected *C*_*α*_ atoms that are perturbed, *δ*(*x*) is a Delta function equal to zero for any nonzero *x*, and unity when *x* = 0. Ω_*j*_ are forces exerted toward 133 directions defined in section 2b. With the full-atom **F** matrix, we further define the residue level CGCM by summing the counts belong to each residue pair such that

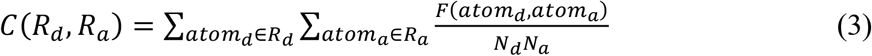

where *R*_*d*_ and *R*_*a*_ are donor and acceptor residues, respectively; *N*_*d*_ and *N*_*a*_ are the number of atoms in the donor and acceptor residue, respectively.

To model the signals propagation process during the hydride transfer reaction, we perturbed 21 sites including *Cα* atoms within the distance of 7Å from the catalytic center (N5 atom of the DHF) as well as four sites locating at cofactor and substrate (see **Fig S1**). Consequently, 133 evenly distributed impulse forces defined in section 2b are applying at each perturbed site; all signals are summed up to one CGCM with elements of *C*(*R*_*d*_, *R*_*a*_) which are then normalized with the 133 directions. In the CGCM, the elements in or immediately near the diagonal, indicating intra-residue communication or that between neighboring residues in primary sequence, have the largest values. The high *C*(*R*_*d*_, *R*_*a*_) score (Communication Scores or CSs in short) along diagonals is intuitive for their strong covalent binding but less interesting in terms of allostery. We pay our attention to the off-diagonal elements satisfying |*index*(*R*_*d*_) *− index*(*R*_*a*_)| *> 2*, which provide information on the signal propagation between long range contacts (say, within or between secondary structures). Several hubs having the highest CGCM scores are the residue pairs that frequently communicate dynamic signals in one or multiple pathways.

**Directionality**: it is worth noting that the pairwise communication in CGCM is asymmetric; that is to say for residue *i* and *j, C*(*R*_*i*_, *R*_*j*_) *≠ C*(*R*_*j*_, *R*_*i*_) despite that the two numbers are usually quite close. As a result, the directionality of signal propagation can be said as from residue *i* to *j* if *C*(*R*_*i*_, *R*_*j*_) *> C*(*R*_*j*_, *R*_*i*_) (see Fig S4 for an example).

For a given residue, a unique “**communication score (CS)***”* can be assigned as the highest CGCM score among pairs formed by this residue and any other non-consecutive residue in the protein – in other words, residue *i*’s CS is the highest score in either the *i*-th row or the *i*-th column (corresponding to donors or acceptors) of CGCM. The CS of each residue is listed in **Table S1**.

### 2e. Evolutionary importance of the ICCs

The evolutionary conservation of residues is compared with their CSs. The sequence conservation of residues are calculated by multiple sequence alignment conducted at the ConSurf web site (66-68), where residues are categorized into 9 conservation levels by “the ConSurf score*”* from 1 (the most diverse) to 9 (the most conserved). We then exam the distribution of the ConSurf score for residues with CS larger than several thresholds - 1.1, 1.2, 1.3 and 1.4.

### 2f. The fold reduction of hydride transfer rates in DHFR mutants is presented as the difference of free energy changes

Suppose that the mutation of an amino acid would break and/or rewire the intrinsic dynamics network, then the chemical modifications (through mutagenesis) in frequently communicating sites, identified by physics approaches (such as td-LRT), could impair the function in the active site allosterically. Here, the forward hydride transfer rate, the chemical step extensively characterized by kinetic isotope experiments (7, 10, 11, 21-32), of wild type DHFR is defined as *k*_*WT*_ and that of its mutant is defined as *k*_*mut*_. Converting the reaction rates to its activation free energy, we can use the inverse Boltzmann relation to obtain Δ*G* = *−RTln*(*k*). Consequently, the difference in activation free energy of hydride transfer reaction between the wild type and a mutant can be written as ΔΔ*G* = *−RTln*(*Γ*), where *Γ* =*k*_*WT*_*/k*_*mut*_ that is the fold reduction of the hydride transfer rate due to a single point mutation. *RT* = *0*.*6* kcal/mol. As can be seen in **Table 1**, a large *Γ* value means that the hydride-transfer rate is largely suppressed by the mutation, while *Γ* near unity means that the reaction rate stays unchanged. The correlation between the CS of each sites and the corresponding averaged change in free energy difference ⟨ΔΔ*G*⟩ is reported in this study, where ⟨*…* ⟩ denotes the average over different mutations at the same site. The results are reported in section 3e.

**Table 1.** Top 6 FDA-approved drugs that could bind ATG4B’s open form via the allosteric site.

| Rank | Drug name | Binding affinity (kcal/mol) | Drug-Site Distance<sup>#</sup> (Å) | Ranking score |
| --- | --- | --- | --- | --- |
| 1 | Fenretinide | -8.2 | 5.0 (6.1) | 1.86 |
| 2 | Moxidectin | -8.0 | 6.3 (6.0) | 1.43 |
| 3 | Conivaptan | -9.0 | 6.7 (7.6) | 1.14 |
| 4 | Lusutrombopag | -7.4 | 6.4 | 0.86 |
| 5 | Halcinonide | -7.3 | 6.3 | 0.86 |
| 6 | Butaclamol | -7.3 | 6.5 | 0.57 |

The drug docking poses are rank-ordered by the average ranking scores considering both the pose binding affinity and distances from poses to the target site (see Methods). Docking is conducted by Vina.

### Drug docking and MD simulations to repurpose FDA approved drugs that target the tdLRT-identified allosteric site of ATG4B

The ATG4B structure adopting an active enzyme conformation (or the “open form*”*; PDB ID:2Z0D; Satoo et al., 2009) was taken for screening of FDA drugs that bind the identified allosteric site. All the missing loops were patched and the catalytically inert mutation, H280A, was back-mutated with the aids of SWISS-MODEL web service (Waterhouse et al., 2018).

Hydrogen atoms of ionizable residues in the ATG4B structure (Satoo et al., 2009) were added or removed per their protonation states, calculated by PDB2PQR web service (Dolinsky et all., 2007). A set of 2016 FDA-approved drugs compiled from the catalogs of MedChemExpress (MCE) FDA-Approved Drug Library (Cat. No.: HY-L022) and Screen-Well® FDA Approved Drug Library (Version 1.5) of Enzo Life Sciences, Inc. were used for the drug screening. The 3D structures of drugs were built by BIOVIA Discovery Studio (Dassault Systèmes BIOVIA, Discovery Studio Modeling Environment, Release 2017, San Diego: Dassault Systèmes, 2016) with appropriate adjustment of the protonation state of all ionizable functional groups under pH = 7. The docking software, AutoDock Vina (Trott et al., 2009), was used to perform virtual drug screening. The PDBQT files required as the input files were generated using AutoDock Tools (Morris et al., 2009). The search box was adjusted to cover the whole protein structure for global search, with a space of 5 Å thickness padded from each side of the box to the protein. The exhaustiveness was set to 100. For each drug, 20 docking poses were allowed to be generated by AutoDock Vina.

To analyze the docking result and choose the drug candidates for repurposing as ATG4B intramolecular allosteric drugs, besides the binding affinity obtained for each docking pose, we also calculate the distance, as a second feature, from the pose to the target sites. Within the top 10 ICCs having the highest communication scores (CSs), we select those >10 Å away from the active sites and spatially clustered to discover a single **allosteric site** comprising TRP27, TYR33, ARG31, LYS39, LYS32 and ILE28 as shown in the red spheres of Fig1b.

We defined the pose distance as the shortest distance between the heavy-atom mass center of a drug and the closest heavy atom of an ICC residue in the allosteric site. To identify relatively high-affinity drugs in short distance from the target site, we retained only the docking poses with affinity *≤* -7 kcal/mol and distance*≤* 7 Å. This resulted in 7 poses from 6 drugs. The poses were then rank-ordered in accord with affinity and distance. The poses are first ranked by affinity only (low affinity ranks higher) and distance only (the shorter, the better) before they are ranked based on the smallest sum of the ranks or the rank-converted scores described in Equation 1^72^ ( see also https://www.biorxiv.org/content/10.1101/2021.01.31.429052 ).

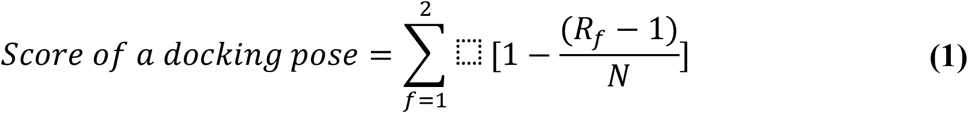

where feature *f* represents the pose affinity or distance. *R*_*f*_ represents the rank of a pose according to the *f* feature (the highest rank is 1; the lowest is *N*). *N* is the total number of the retained poses, which is 7 in this case.

Following standard protocol described in DRDOCK^72^, we have performed 10 ns MD simulations for the top 3 drug candidates in Table 1. The last 500 ps trajectory is used to calculate the MM/GBSA-derived affinity and contact number.

### Protein expression and purification

Human ATG4B with N-terminal 6xHis-tag and human LC3B with both N-terminal 6xHis-tag and C-terminal S-tag were cloned into pETDuet-1 expression plasmids as previously described *(Shu, CW et al*., *Autophagy 6:7, 936-947, 2010)*, subsequently transformed into *E. coli* BL21 (ECOS 21, Yeastern Biotech Co., FYE207-40VL). A single colony harboring the correctly in-frame inserted gene was inoculated into an overnight culture that was then diluted 100-fold in fresh Luria-Bertani broth with 50 µg/mL ampicillin growing at 37°C with 200 rpm agitation for 2 to 3 hrs until the optical density of the culture at 600 nm reaches 0.4 (ca. 3 × 10^8^ cells/ml). The recombinant proteins were then induced by 0.1 mM Isopropyl β-D-1-thiogalactopyranoside (IPTG) and cultivated at 20°C for another 5 hrs.

The expression cell culture was harvested by 12,000×g centrifugation at 4°C for 20 mins and the cell pellets of 1 liter broth were resuspended in a 10 mL lysis buffer (50 mM Tris-HCl, pH7.4, 300 mM NaCl, 20 mM imidazole). The 10-mL cells were lysed by sonication and centrifuged at 38,500×g for 15 mins at 4°C and the supernatant containing the 6xHis-tagged recombinant proteins was filtered by 0.8 μm membrane before subjected to 3-mL Ni-NTA-agarose (Qiagen, 30250) gravity column. Subsequently, sample buffer comprising 10 mM Tris-HCl, pH 7.4, 150 mM NaCl, 10 mM β-mercaptoethanol, 0.1% triton X-100 was applied throughout the purification, including pre-washed by 20 mM imidazole and collected elution with 100 mM imidazole. The ATG4B and LC3B proteins were concentrated and frozen with 20% glycerol. The protein purity was verified >90% by coomassie blue staining on SDS-PAGE and the quantity was measured by bicinchoninic acid assay.

### ATG4B enzyme activity assay by S-tag-based immunoblotting

ATG4B enzyme activity assay was analyzed by immunoblotting. The purified recombinant ATG4B (5-10 nM) was incubated with 500 nM C-terminal S-tagged LC3B in 100 µL reaction buffer containing 50 mM Tris-HCl, pH 8.0, 150 mM NaCl, and 1 mM dithiothreitol at 37°C for 2 hrs. Reactions were stopped by addition of 5X SDS-sample buffer and 95°C heated for 5 mins before loaded to 12% SDS-PAGE gel for electrophoresis, after which the samples were then transferred to nitrocellulose membranes (PALL, Biotrace NT 66485) for immunoblotting analyses. The membrane was blocked with 5% skim milk (Sigma-Aldrich, 70166) in TBST buffer (TBS with 0.05% Tween-20) for 1 hr at room temperature with mild shaking, and then incubation with anti-S-tag (Bethyl, A190-135A), anti-c-Myc (Sigma-Aldrich, C3956) or anti-ATG4B (Sigma-Aldrich, A2981) primary antibodies in TBST buffer containing 5% BSA for overnight at 4°C with mild shaking. The proteins were then probed with peroxidase conjugated mouse anti-rabbit IgG secondary antibody (Santa Cruz, sc-2357-CM) with 5% skim milk in TBST buffer for 1 hr at room temperature. The membranes were then treated with enhanced chemiluminescence (ECL) reagent (GE Healthcare, RPN2232) for band intensity detection by the ImageQuant™ LAS 4000 Imaging system (Cytiva, USA). The intensities of remaining substrate LC3B after enzymatic cleavage were quantified by the software ImageJ.

### Tumor cell culture and cell viability assay by WST-1

Three tumor cells were tested in this study - HCT116 (colorectal cancer), AsPC-1 (pancreatic cancer) and MDA-MB-468 cell lines (breast cancer). Human colorectal carcinoma cell line HCT 116 (catalog number: BCRC 60349, Bioresource Collection and Research Center (BCRC), Hsinchu City, Taiwan) was maintained in the culture medium: McCoy’s 5a medium (catalog number: 16-600-082, Gibco, Waltham, MA) with 1.5 mM L-glutamine (Gibco) and 10% fetal bovine serum (FBS, Gibco). Human pancreatic adenocarcinoma cell line AsPC-1 (catalog number: BCRC 60494, BCRC) was maintained in the culture medium: RPMI 1640 medium (catalog number: 11875168, Gibco) with 2 mM L-glutamine, 4.5 g/L glucose (Gibco), 10 mM HEPES (Gibco) and 1 mM sodium pyruvate (Gibco) and 10% fetal bovine serum. Human breast adenocarcinoma cell line MDA-MB-468 (catalog number: ATCC® HTB-132™, American Type Culture Collection (ATCC), Manassas, VA) was maintained in the culture medium: Leibovitz’s L-15 medium (catalog number: 11415064, Invitrogen, Waltham, MA) with 2 mM L-glutamine and 10% fetal bovine serum. All cells were incubated in a humidified atmosphere without CO_2_ in air at 37°C to serve as the control for the following assays.

For cell viability assay, tumor cells (1 × 10^5^/mL) were suspended in the medium without testing drugs and then inoculated in 96-well plates (100 μl per well) for attachment first. After a 12-hour incubation to allow for cell attachment, the conditioned medium was withdrawn. Then, Fenretinide and Moxidectin (10 μM) were added to the culture medium (100 μl per well) to treat cells for two additional days. Cell proliferation was quantified by performing a Premix WST-1 Cell Proliferation Assay System (Takara, Japan). According to the manufacturer’s protocol, cell proliferation was determined by measuring the optical density (OD) after the reaction for 3 hours by recording the absorbance at 450 nm using a plate reader (Multiskan Go, Thermo Fisher Scientific).

### Statistical analysis

Cell culture and Biochemical data are expressed as the mean ± SEM from 3 individual experiments. A nonparametric two-tailed Student’s t-test are performed for our analysis with the P values less than 0.05 being considered significant.

## RESULTS

### The signal propagation and characteristic time

Movie S1 and Fig. 2(a) show the relaxation motion of atom *i* (the *C*_*α*_ atom of residue G121) as a function of time, where the point impulse force acting on the atom *j*, the N5 atom of the DHF at the catalytic center, toward the z-direction, is denoted by (*θ*_*j*_, *ϕ*_*j*_) = (*0,0*); the characteristic time, 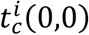, is 2.6 ps. Considering the force ensemble act at the catalytic center, the red circle in Fig. 2(b) shows the average characteristic time of every single atom *I* over the force ensemble, 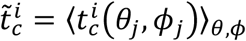, and the black bars denote the corresponding standard deviation. It can be seen that the signal starting from the catalytic center can almost go through the entire DHFR within few picoseconds. Further, 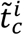 as a function of the distance between atom *i* and *j, d*_*ij*_, is shown in **Fig. 2c**. The linear regression of 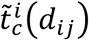 marked by blue line gives a propagation speed of ∼580 *m/s* with a correlation of ∼0.8. Generally, the data show that the atom *i* with larger 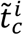 is farther in distance from the perturbed site and has larger variance. The variances suggest that the speed of signal is not isotropic and could be affected by the intramolecular dynamic network of residues.

### The Longest Dissemination Time (LDT) and disseminators

The dynamic network inside a protein also regulates how fast a perturbation could propagate through the entire protein. In this study we perturb the C_*α*_ of all the 159 residues in DHFR, also the C6 atom of the NADPH and N5 atom of the DHF. The LDT of four groups in NADPH and three groups in DHF are also noted (**Fig S2**). For each residue *j*, a perturbation force ensemble in 133 different directions is used and then the slowest 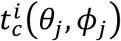at a given residue *i* is noted as the longest dissemination time (LDT) of the residue *j*. The residues with short LDT are called “disseminators*”* (54). Figure 3 shows the LDT for all the residues. It was found that several active sites (ILE 5, Phe 31, ILE 94, C6 of NADPH and N5 of DHF) are among good disseminators. The results suggest that the active sites locate at the optimal position of network to efficiently broadcast the mechanical signals.

**Figure 3.**
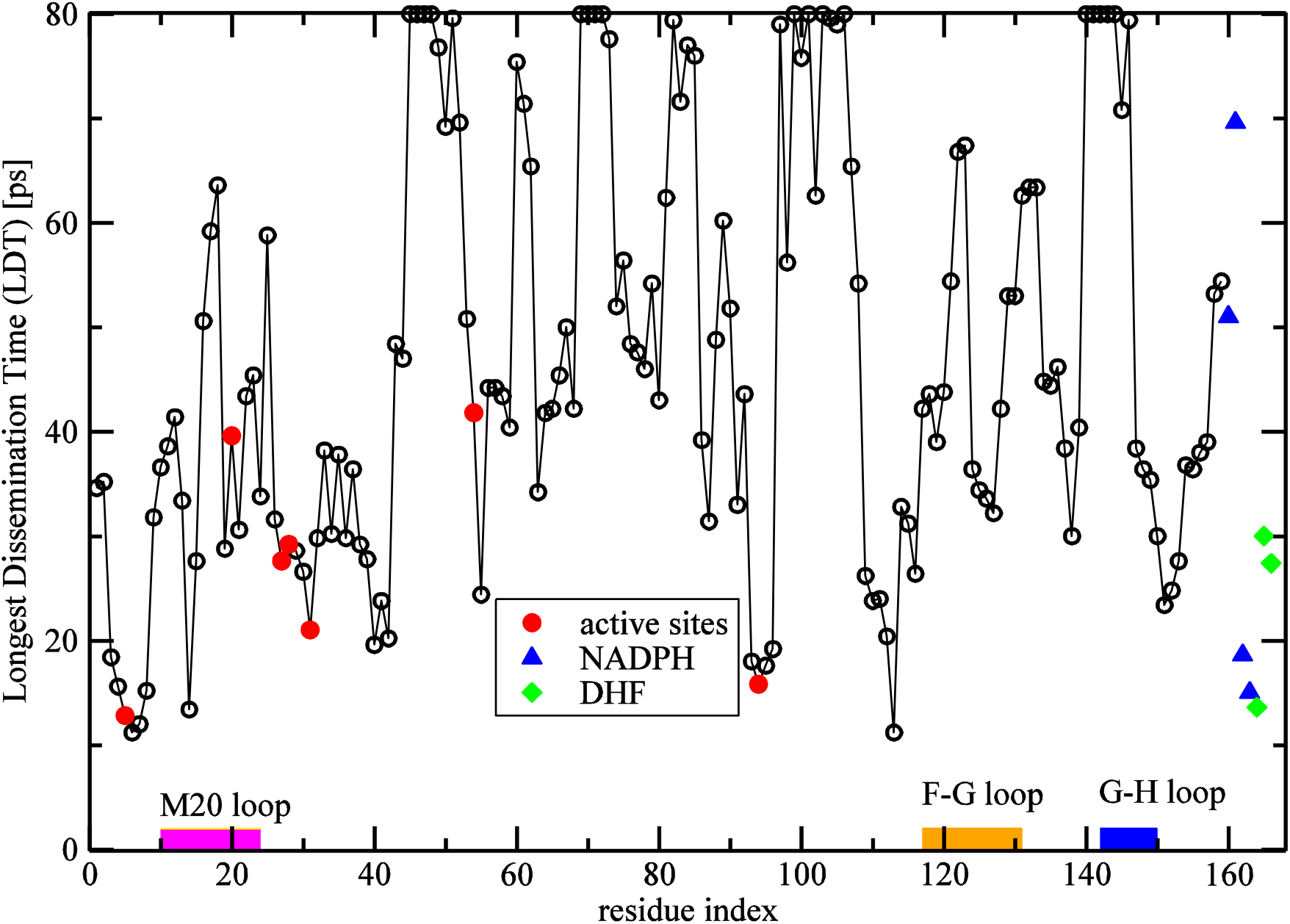
The longest dissemination times (LDTs) (54) of selected indices of sites. An LDT is the slowest response time defined by the longest *t*_)_ in the DHFR with 133 evenly distributed impulse forces starting from a *C*_*α*_ atom of a selected residue. Indices 160-163 (in blue triangle) and 164-166 (in green diamond) indicate 4 and 3 groups of NADPH and DHF, respectively (see Figure S2). Index 163 is C4 of nicotinamide group of NADPH and index 164 is C6 of pterin group of DHF; both 163 and 164 are good disseminators with the smallest LDTs among all the perturbation sites (residues). For our studying purpose, we do not concern large LDTs; hence an estimation of LDT longer than 80 ps is shown as 80 ps. The solid red circles indicate the active sites: Ile5, Met20, Asp27, Leu28, Phe31, Leu54 and Ile94 (62) showing relatively small LDT among all.

### 3c. The Coarse-Grained Connection Matrix (CGCM) and communication score (CS)

Following the procedure in section 2d, the CGCM of DHFR can be obtained (**Figure 4a**). There are several hot spots in the off-diagonal elements, especially, the largest two elements are found to locate at C(13,122)=1.52 and C(13, 121)=1.42 - the donor-acceptor pairs of V13-D122 and V13-G121, respectively. Previous experiments have shown that the mutations in D122 and G121 can significant obstruct the hydride transfer rate. Interestingly, as shown in **Table S3**, we decomposed C(13,121) into 21 components by the perturbed sites, and it is found that signals from the perturbed C6 of the pterin group (site 164) and Y100 are almost twice the size of signals from other perturbed sites. The results provide an evident supporting that a strong coupling between the residue G121 and the catalytic center. Consequently, the CSs are determined following section 2d. **Figure 4b** shows the color-coded DHFR structure by **the CSs**, where the lowest and highest CS correspond to dark blue and dark red, respectively. The first few highest CSs can be found at the residue D122 as well as the second highest CS at the residue G121 located in the F-G loop. Some other residues such like M42, L54 and D27 show moderate CS of 0.6-0.7, while G67, S148 and W133 show CS lower or equal to 0.5. With the current method where pairwise communication is not symmetric, we can assign the directionality for the signal flow. For instance, we can see how the signal is propagated from the allosteric site at G121 and D122 to residues near the catalytic center (**Fig S4**).

**Figure 4.**
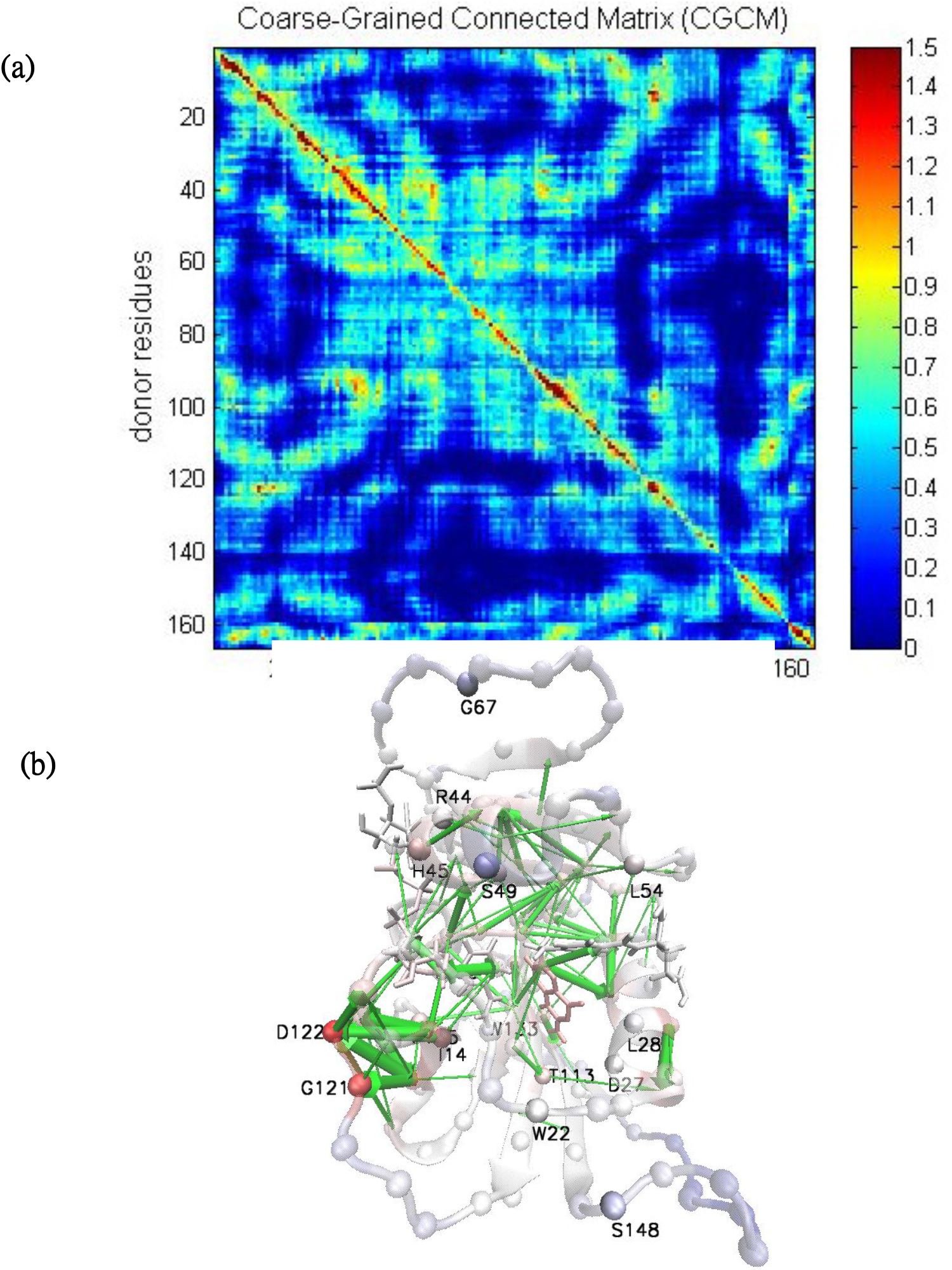
**(a)** The coarse-grained connection matrix (CGCM) in which each element, *C*(*R*_*d*_, *R*_*a*_), records the counts of mechanical signals propagating from a donor (*R*_*d*_) residue to an acceptor (*R*_*a*_) residue, generated by the method described in section 2d. Consequently, a unique “communication score (CS)*”* is assigned to a selected residue with the highest CGCM off-diagonal element satisfying |index(*R*_*d*_)-index(*R*_*a*_)| > 2 pertaining to that residue (either a donor or an acceptor). In other words, the CS of residue *i* is the highest CGCM score of the *i-th* row or the *i-th* column. For example, selecting the residue 122, as a donor, the highest CGCM off-diagonal elements in the 122-th row is *C*(1*22*,1*3*) = 1.39; as an acceptor, the highest CGCM in the 122-th column is *C*(13,122) = 1.*52*, therefore, the CS of residue 122 is 1.52. **(b)** The CSs color-coded 3D structure of the DHFR is plotted by the software VMD, where cartoon and spheres (*C*_*α*_ atoms) are colored by mapping the minimal CS (dark blue) and the maximal CS to unity (dark red). The green arrows pointing from donors to acceptors indicate signal propagation pathways, where the thickness represents the size of CS for the pair of interest.

After collecting mutagenesis data from 35 mutants at 17 sites of which the hydride transfer rates were carefully characterized without blending in kinetics of substrate binding or product release (7, 10, 11, 21-26, 28), the ratio (*Γ*) of wide-type hydride transfer rate (7, 10, 11, 21-26, 28) to that of a single mutant can be obtained. It is found that the hydride transfer rate of DHFR could drastically fall by 2 or 3 orders (resulting in a *Γ* value of hundreds or more than a thousand; see Table 1) if a residue with high CS is mutated. It happens even when the residue is remote (>15Å) to the catalytic center (**Table 1, Fig 6**), suggesting that allosteric regulation on activity of a functional site can be imposed by perturbing (chemically herein, by mutagenesis) spatially clustered ICCs (e.g. G121 and D122) with a high CS, if medicinal interest of this study is concerned.

**Figure 5.**
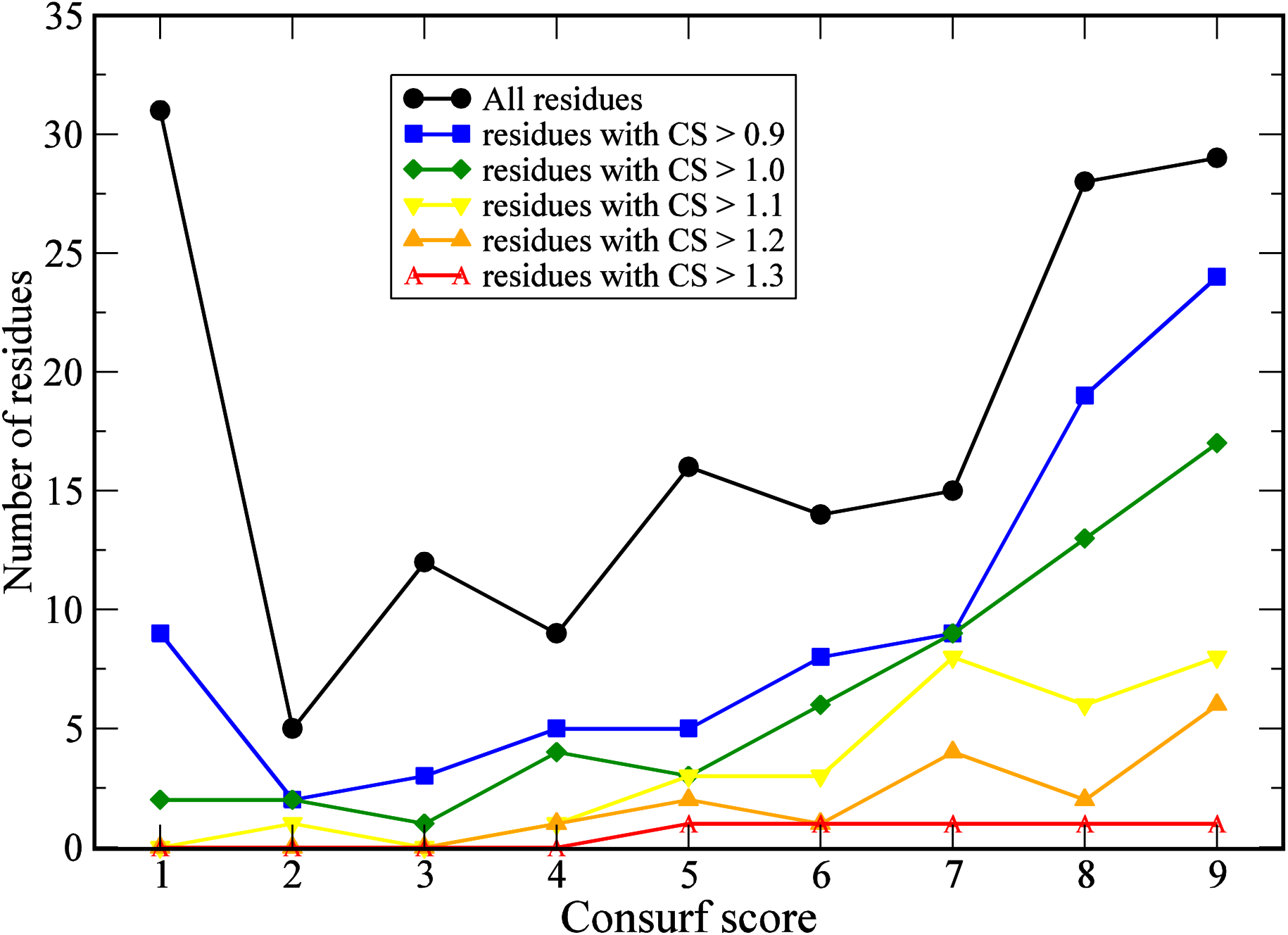
The histograms of conservation scores (66-68) for the residues of different CSs, where score 9 indicates the highest evolutionary conservation and 1 the lowest based on multiple sequence alignment results (66-68). The black line indicates the distribution for all 159 residues of DHFR. The blue, green, yellow, orange and red lines correspond to the residues with CSs higher than 0.9, 1.0, 1.1, 1.2 and 1.3, respectively.

**Figure 6.**
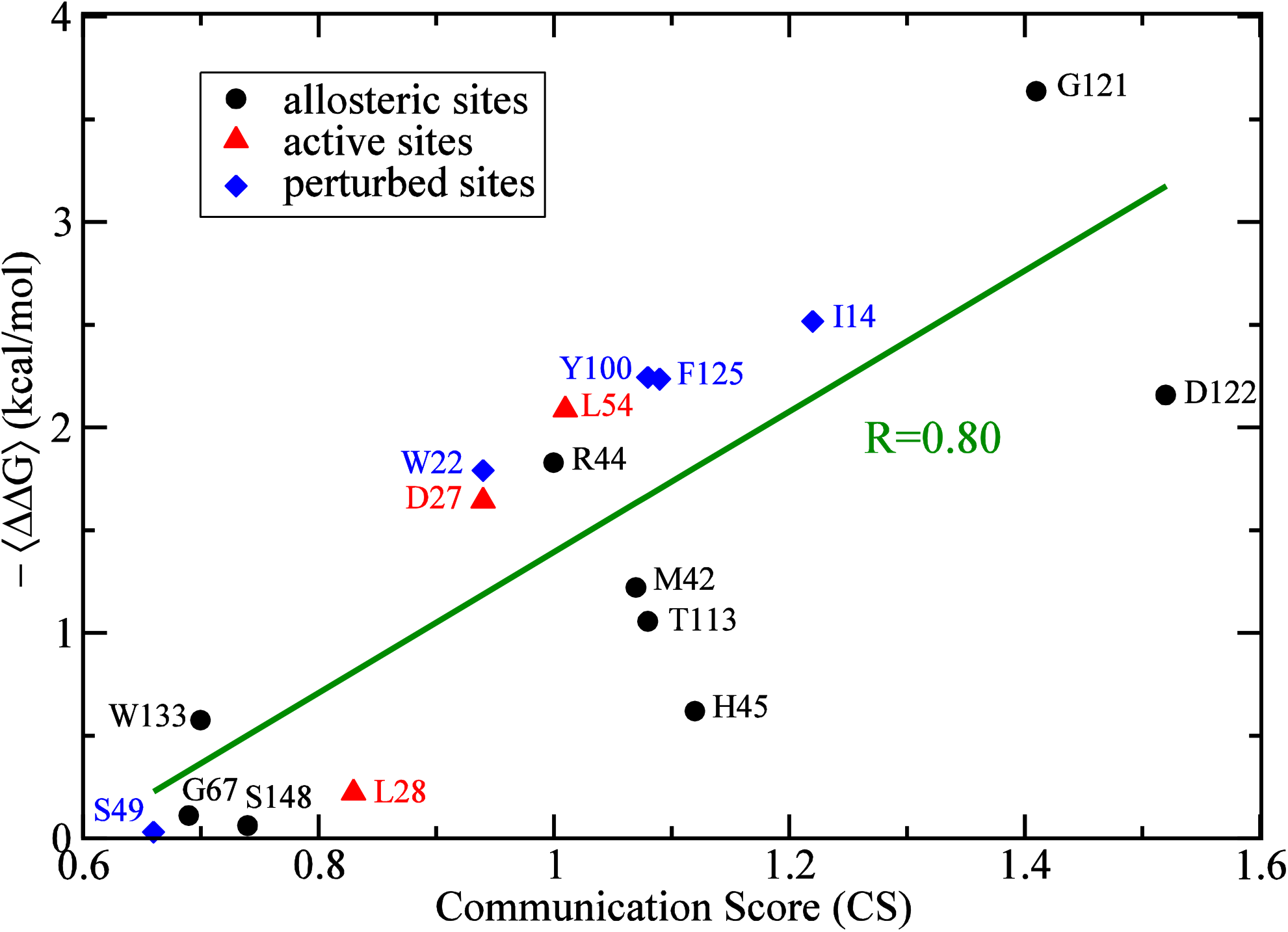
The averaged change in free energy difference of the hydride transfer rates between the wild type and its mutants is plotted against the communication score (CS). The averaged ⟨ΔΔ***G***⟩ = ⟨Δ*G*_*WT*_ *−* Δ*G*_*mutant*_⟩ = ⟨*−RTln*(*k*_*WT*_*/k*_*mut*_)⟩, where *k*_*WT*_ and *k*_*mut*_ are the hydride transfer rates of wild type DHFR and its mutant, respectively. ⟨*…* ⟩ denotes the average over different mutants at the same site; the data are listed in **Table 1**. The CS for each residue is determined following the method in section 2d, and CSs for all the residues in DHFR are listed in **Table S1**. The red triangles are active sites, blue diamonds are perturbed sites and black circles are allosteric sites (>15Å from the catalytic center). The dark green line shows linear regression of the data for 17 sites with a correlation coefficient of 0.80.

### The conservation of residues of DHFR in evolution

We further examined the evolutional importance of the identified ICCs. The number histogram of the ConSurf scores for all residues in DHFR is shown in **Figure 5** (the black line). There are 31 residues with the lowest ConSurf score (that is 1) as well as 57 residues with the highest ConSurf scores (8 or 9) which dominate the population. As examining the histograms of ConSurf scores for residues having a CS >0.9 (84 residues; in blue line), >1.0 (57 residues), >1.1 (30 residues), >1.2 (16 residues) and >1.3 (5 residues; in red line), we found the population gradually shifts toward evolutionarily conserved regions. The finding supports the aspect that dynamically significant residues are also evolutionarily conserved (69).

### The exponential dependence between the hydride transfer rate reduction and CS

By collecting experimental data from the literature, as shown in the **Table 1**, we summarized the fold reduction of hydride transfer rate in 35 mutants at 17 sites. The fold of hydride transfer rate reduction for a point mutation is expressed as *Γ* = *k*_*WT*_*/k*_*mut*_. For example, the I14A mutation causes 40 fold of hydride transfer rate reduction. The red and blue highlights indicate active sites and perturbed sites, respectively. ⟨ΔΔG⟩ = ⟨ΔG_*WT*_ *−* ΔG_*mutant*_⟩ = ⟨*−RTln*(*Γ*)⟩ is the average for a set of difference of activation free energy in the hydride transfer rates between the wild type and its mutants at a given site (average over the rows in **Table 1**), where ⟨*…* ⟩ denotes the average of ΔΔG for mutants at the same site. **Figure 6** reveals there is an appealing correlation (>0.8) between mutants’ -⟨ΔΔ*G*⟩ and their communication scores, suggesting that the frequency of dynamic signal propagation through a residue could play an important role in allosteric control over the hydride transfer rate.

### A tdLRT-revealed intramolecular allosteric site of ATG4B was targeted to confer tumor suppression synergy

To further validate this td-LRT-based method, we would like to take a disease target protein whose intramolecular allosteric site(s) has not been previously reported. The autophagic protein ATG4B, an cysteine protease involved in the regulation of autophagosome formation and previously shown to drive the oncogenic process in progressed tumors^70, 71^, is chosen for our purpose. We aim to first predict the clustered intramolecular communication centers (ICCs) as potential allosteric sites, in a notable distance away from the catalytic center, and then find the drugs that can bind the sites. We screened a library of 2016 FDA-approved drugs^72^ by small molecule docking for this purpose. These drugs are then tested for their *in vitro* inhibition of ATG4B’s function by enzymatic assay and suppression of tumor cell growth.

By perturbing 41 residues within 8Å from Cys74, the catalytic cysteine of ATG4B, we were able to use the same approach as we did for DHFR to derive the coarse-grained communication map (CGCM)(Fig 1a) where a group of “hot-spots*”* near the residue 30 in the off-diagonals of CGCM can be found. Among the top 10 ICCs, 6 are spatially close enough to be clustered into a single site, 26.1Å away from the catalytic center, as shown in Figure 1a,b. Our docking study (see Methods) found, from 2016 FDA drugs, only 6 FDA drugs having relatively high affinity (< -7 kcal/mol) and site proximity (<7Å) that could bind the site (Table 1). In the 6, Fenretinide, Moxidectin and Conivaptan are among the strongest three to bind the allosteric site in ATG4B’s active (open) form (PDB ID:2Z0D), considering both the binding affinity and their pose-site distances (Table 1; also see Method for definition). We also used the closed form of ATG4B (PDB ID: 2CY7) to conduct the same screening procedure. The results show that Fenretinide, Moxidectin and Conivaptan can still be found among the 127 drugs (see SI) that have relatively high affinity (< -7 kcal/mol) and site proximity (<7Å) against the closed form, but not the other three drugs in Table 1. We therefore conduct MD simulations for these three drugs to examine their binding stability and site proximity. The top two results based on the heavy-atom contact and proximity to the allosteric site (Fig 7c) are taken for further enzymatic assays and tumor suppression tests. In the two, Moxidectin has a marginally higher contact (41 vs 38) and closer site proximity (6.0Å vs 6.1Å) than Fenretinide. It is also found to inhibit ATG4B catalysis by 36% and >80% at 10 and 20uM, respectively (Fig 7d), while suppresses tumor growth better in the pancreatic and colorectal cancer cell lines than Fenretinide that seems to be a better breast cancer inhibitor (Fig 7e).

**Figure 7.**
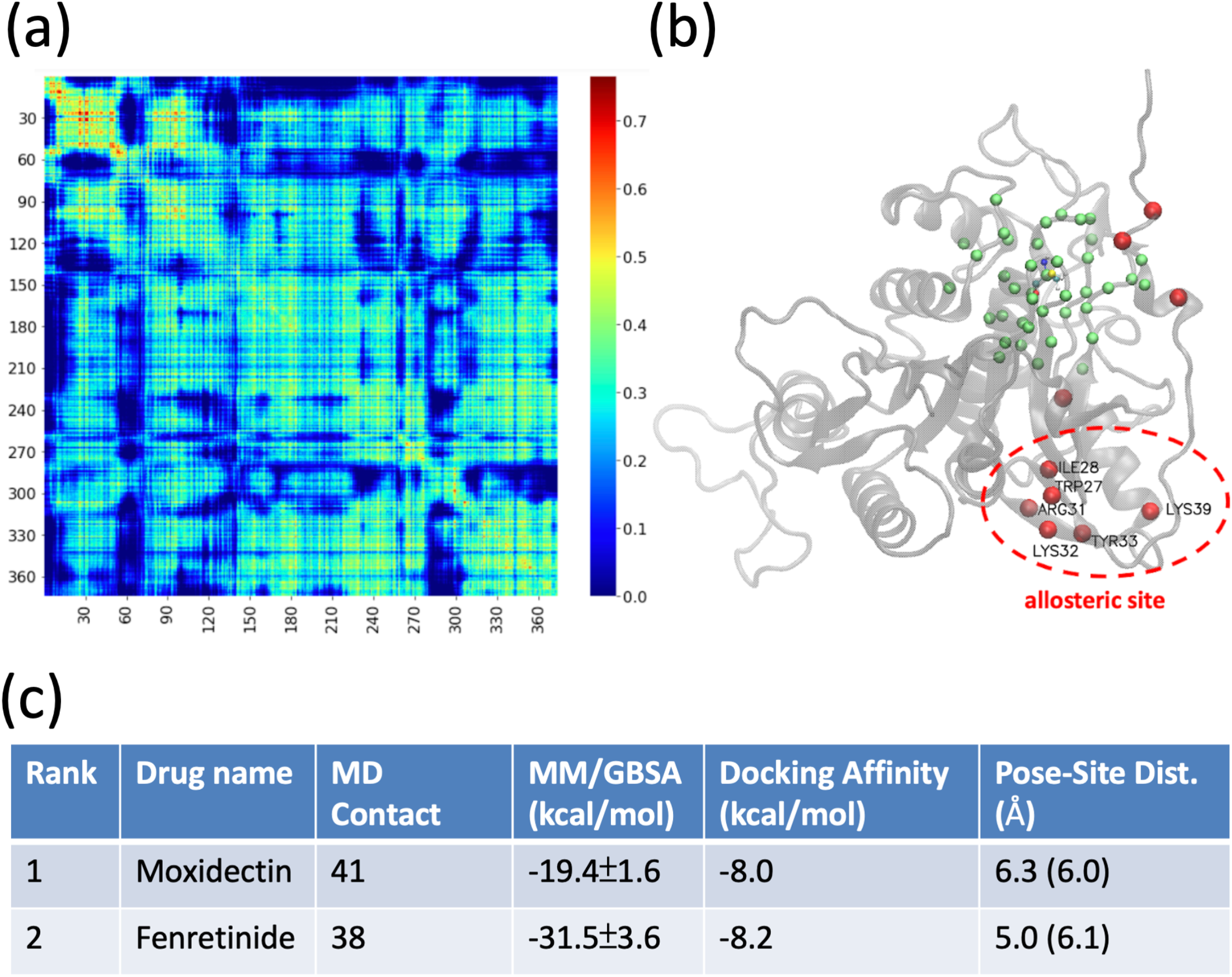
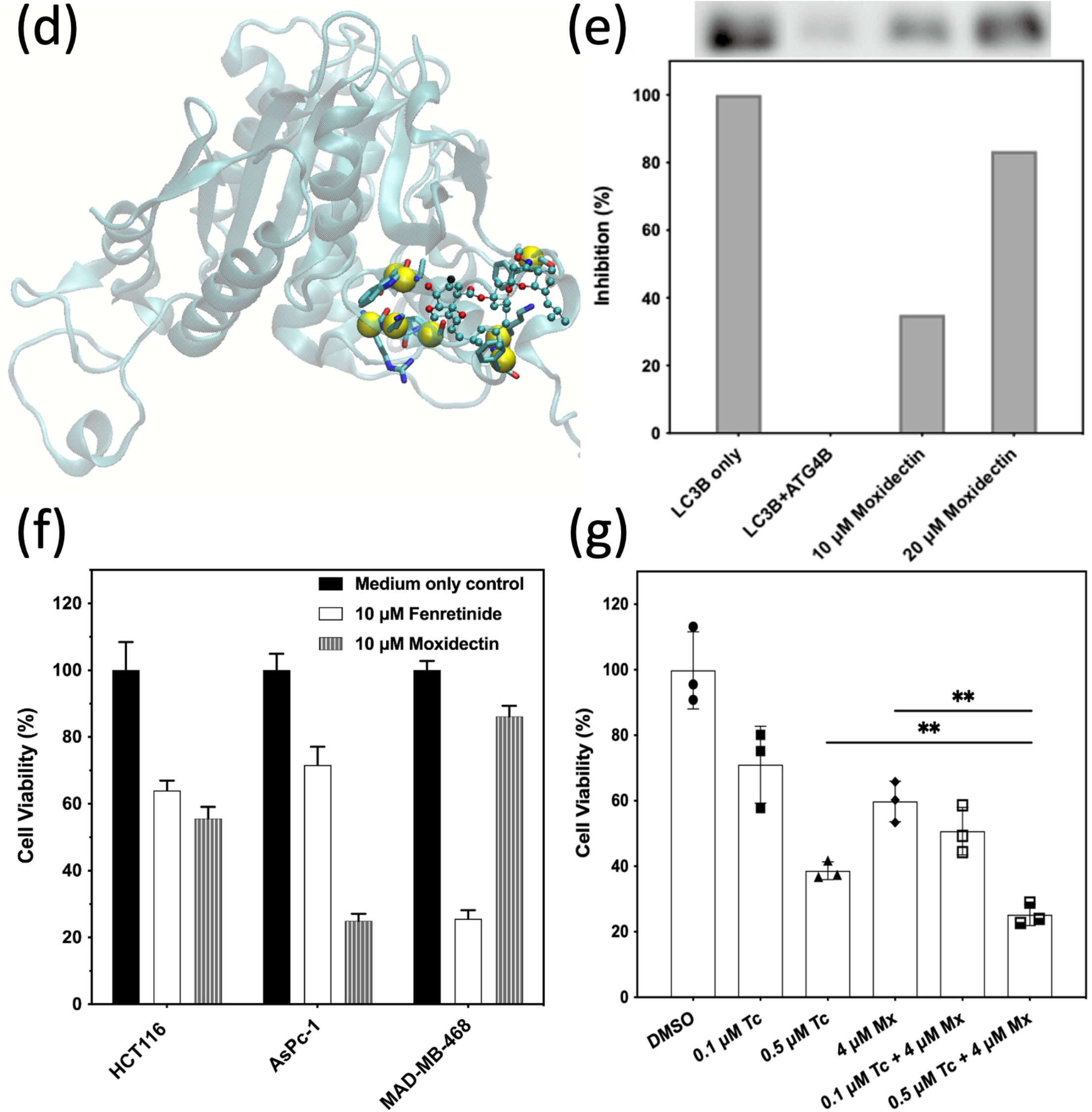
Coarse-Grained Connection Matrix (CGCM) for ATG4B and the clustered ICCs suggesting an allosteric sites. (a) The coarse-grained communication matrix is the average over perturbing the 41 C*α* atom within 8 Å of the Cys74. (b)The td-LRT-derived top 10 ICCs (in red spheres) and their corresponding communication scores in ATG4B are (TRP27, 0.71), (TYR33, 0.71), (ARG31, 0.70), (LYS39, 0.7), (LYS32, 0.69), (ILE28, 0.67), (ARG49, 0.67), (LEU6, 0.67), (ARG12, 0.67), (TYR8, 0.67), where TRP27, TYR33, ARG31, LYS39, LYS32 and ILE28 are spatially clustered as an allosteric site. The average distance between the C*α* atoms of these 6 residues and that of Cys74 is 26.1 Å. **(c)** The top two docking results are subject to further MD analyses including MM/GBSA-based affinity evaluation and pose-site distances characterization; in the Pose-Site distance column, the distances from docking and after 10ns simulation are outside and inside the parentheses, respectively. **(d)** A representative snapshot of how Moxidectin (Mx; in ball-and-stick) interacts with the allosteric residues (in yellow sphere) that are within 4Å of the Mx. **(e)** Inhibition% is defined as (S-tag-blotting intensity of the sample minus that of drug-free ATG4B+LC3B mixture) / (S-tag-blotting intensity of the LC3B only minus that of drug-free ATG4B+LC3B mixture). The higher the inhibition percentage, the better the drug. **(f)** HCT116 colorectal, AsPc-1 pancreatic and MAD-MB-468 breast cancer cell lines are suppressed by 10 μM of the top 2 FDA drugs screened from small-molecule docking and MD simulations. **(g)** 4 μM Mx co-used with 500nM tioconazole, an ATG4B active site inhibitor (Liu et al., 2018), shows an improved tumor suppression efficacy as compared with individual treatment alone (*p*-value < 0.01, Student’s t-test, two tailed).

## DISCUSSIONS

### Is communication frequency between ICCs a function of geometric or physiochemical properties of a protein?

Although the correlation between the allosteric sites and the ICCs is established in our study, the question still remains as for how this “communication*”* property is related to known geometric or physiochemical features in an enzyme. This is not immediately clear at the moment. Our ENM analysis of the DHFR structure showed that the frequent communicators V13 and G121 are not located in the hinge of the slowest mode (suggesting functional importance (67-71)), nor are they located at the peaks of highest frequency modes (indicating the possibility to serve as a local folding core (72)) (Fig S5). The H-bonds formed between V13 and G121 are nowhere standing out as compared to those formed between other pairwise interactions. Thus far we have not found that the communication frequency of ICCs is correlated with other basic geometric or physiochemical properties such as mechanical hinges (68), binding interface (69-71) or high local packing density (72). Yet, they are evolutionarily conserved and regulatory of enzyme catalysis even when situated remotely.

### The nonadditive of the changes of free energy difference for double mutant relating to the coupling of two distal residues also revealed by the elements of CGCM

The coupling between two distal residues has been studied by measuring the changes in free energy difference (ΔΔG = *k*_*A*_*Tln*(*Γ*) where *Γ* = *k*_*WT*_*/k*_*mut*_) for DHFR’s hydride transfer rates between two single mutants and a double mutant of the same two residues (11, 28).

It was proposed that the degree of coupling of two distal residues can be measured by the non-additivity of coupling energy ΔG_*I*_ (11), where the coupling energy Δ_*3*_G_*I*_ between two residue A and B is Δ_*3*_G_*I*_ = ΔΔG_*A,B*_ *−* (ΔΔG_*A*_ + ΔΔG_*B*_). The larger the magnitude of |Δ_*3*_G_*I*_|, the higher degree of coupling between the two selected residues is. For instance, given *Γ* for three single mutants M42W(*Γ* = 41)(11), G121V(*Γ* = 1*66*)(11) and F125M(*Γ* = 4*2*)(28), the coupling energies |Δ_*3*_G_*I*_| of three double mutants: M42W-G121V (*Γ* = *7600*), M42W-F125M (*Γ* = *5*1*2*) and G121V-F125M (*Γ* = *6*1)(28) are 0.077, 0.73 and 2.83, respectively, suggesting positions 42 and 121 are relatively independent, while the 121-125 pair are the most coupled among the three pairs. Interestingly, we found that the communication scores [C(42,121) = 0.26 and C(121,42) = 0.24] are indeed the smallest as compared to [C(42,125) = 0.46 and C(125,42) = 0.45] and [C(121,125) = 0.59 and C(125,121) = 0.66] that is the most frequently communicating pair among the three. Our results show the same order of coupling strength G121V-F125M > M42-F125M > M42W-G121V as that derived from experimentally characterized hydride transfer rates, suggesting the count of propagating signals between a pair of residues can be viewed as a measure to quantify functionally relevant residue-residue coupling in proteins.

### The robustness of CS

We have tested several combinations of perturbation sites involving including or excluding p6, p7, p24 to p159. Overall, the conservation dependence (Fig 5) barely changed, and the correlation between CS and hydride transfer rate reduction in mutants varied slightly (Fig 5a, 5b). In addition to the local structure dynamics such as the flexibility predicted by NMR (8,20,33) or contact number from the structural information, the herein reported CSs could provide a quantitative measurement describing “the propensity of signal propagation between residues*”*. According to the evidence that CSs highly correlate with the hydride transfer rate reduction, our analysis support the aspect that the signal propagation regulates (if not facilitates) the enzyme catalysis, providing an alternative theoretical basis (apart from NMR (8,20,33), bioinformatics, MD) for enzymologists’ further exploration.

**Table 1.**
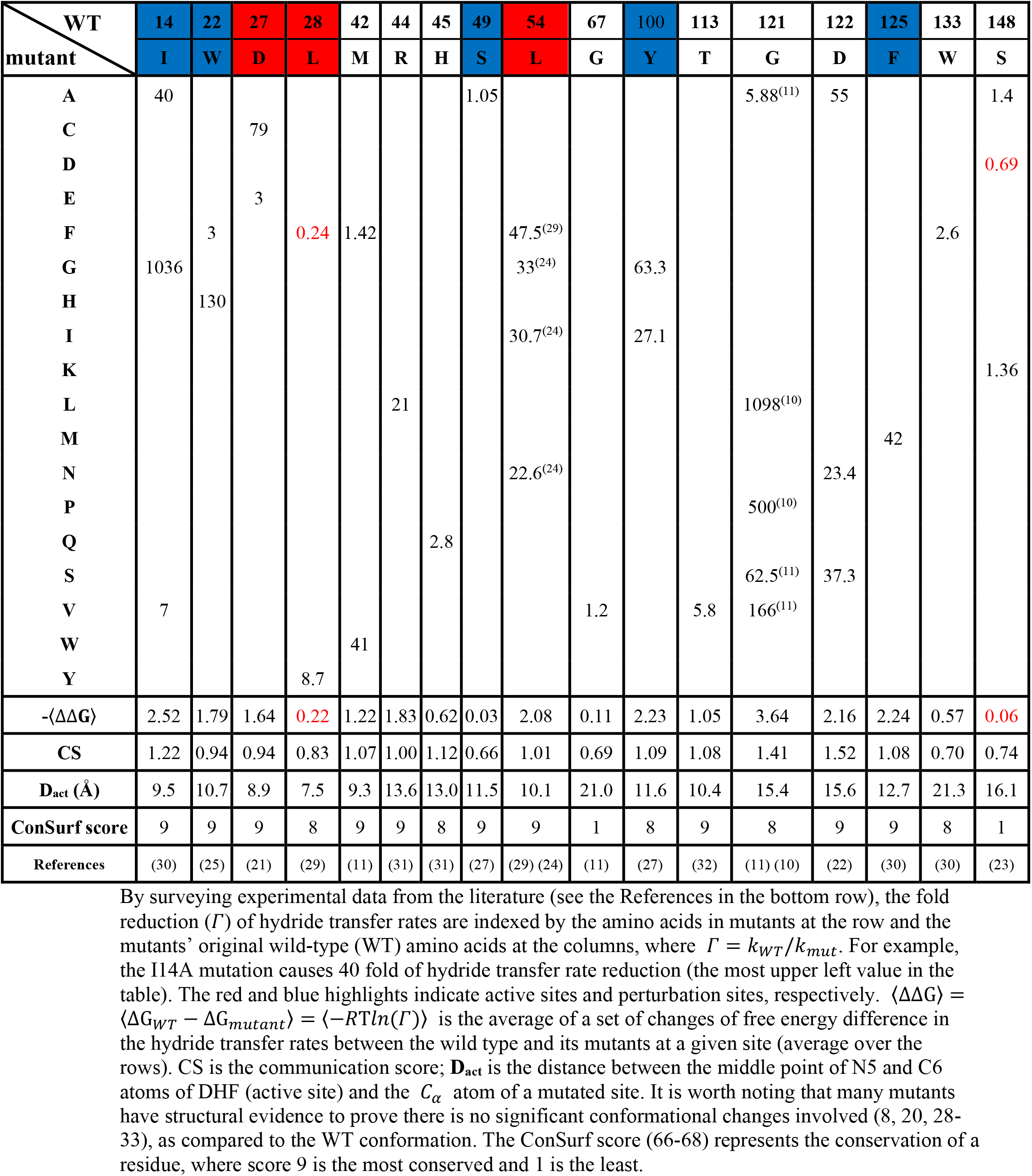
Fold reduction of hydride transfer rates, *k*_*WT*_*/k*_*mut*_, for mutants

## ACKNOWLEDGEMENT

This work was financially supported by National Science and Technology Council of Taiwan (104-2113-M-007-019-). This work used the supercomputers TSUBAME3.0 provided by Tokyo Institute of Technology through the HPCI System Research Project (Project ID: hp190131) in Japan as well as Taiwania I and II provided by National Center for High-performance Computing (NCHC) of National Applied Research Laboratories (NARLabs) in Taiwan.

## SUPPORTING INFORMATION

### Movie of ICCs of DHFR

**The active sites and perturbed sites**

**Figure S1.**
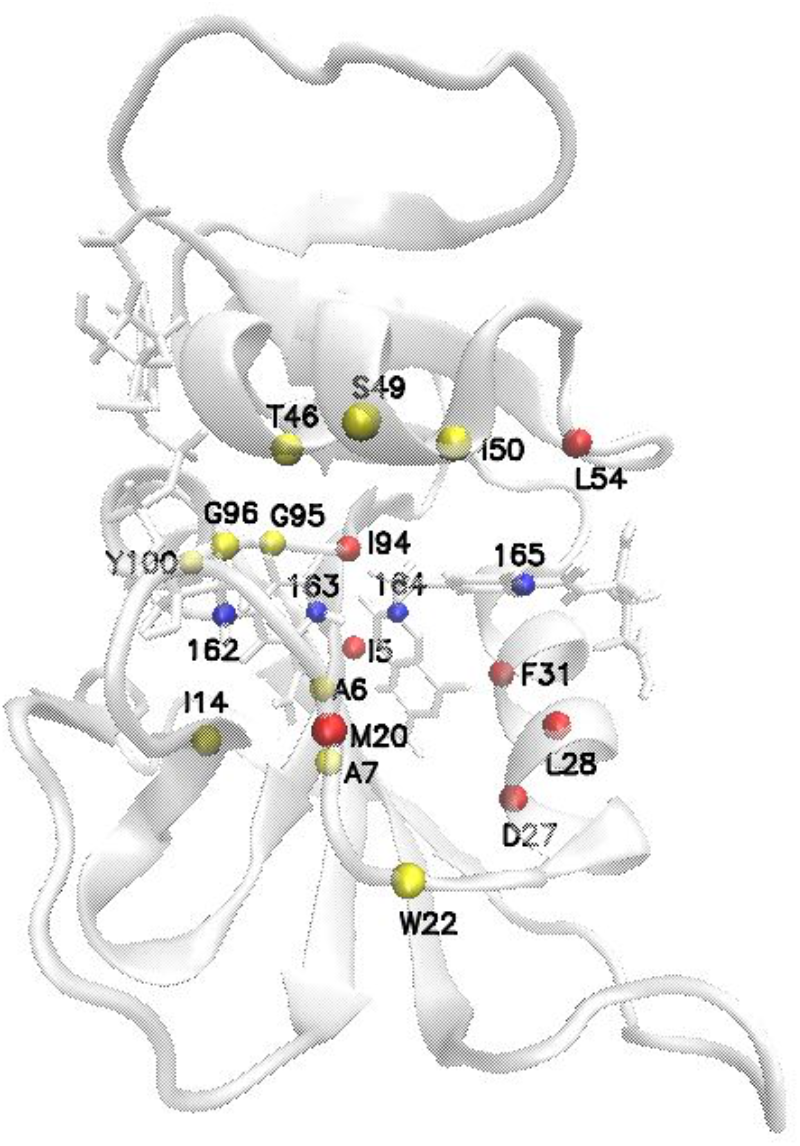
The 21 spheres are perturbed by impulse forces, where the red spheres are active sites: ILE5, MET20, ASP27, LEU28, PHE31, LEU54 and ILE94; yellow spheres are residues within 7 **Å** from the catalytic center: ALA6, ALA7, ILE14, TRP22, THR46, SER49, ILE50, GLY95, GLY96, TYR100. The four blue spheres are the perturbed sites at cofactor and substrate, which is the first Carbon (site 162) of the NADPH at the ribose close to the nicotinamide, the C4 (site 163) of the NADPH at the nicotinamide close to the donor hydride, the C6 (site 164) of the pterin group of the DHF and C15 (site 165) of the benzoyl group of the DHF.

**Figure S2.**
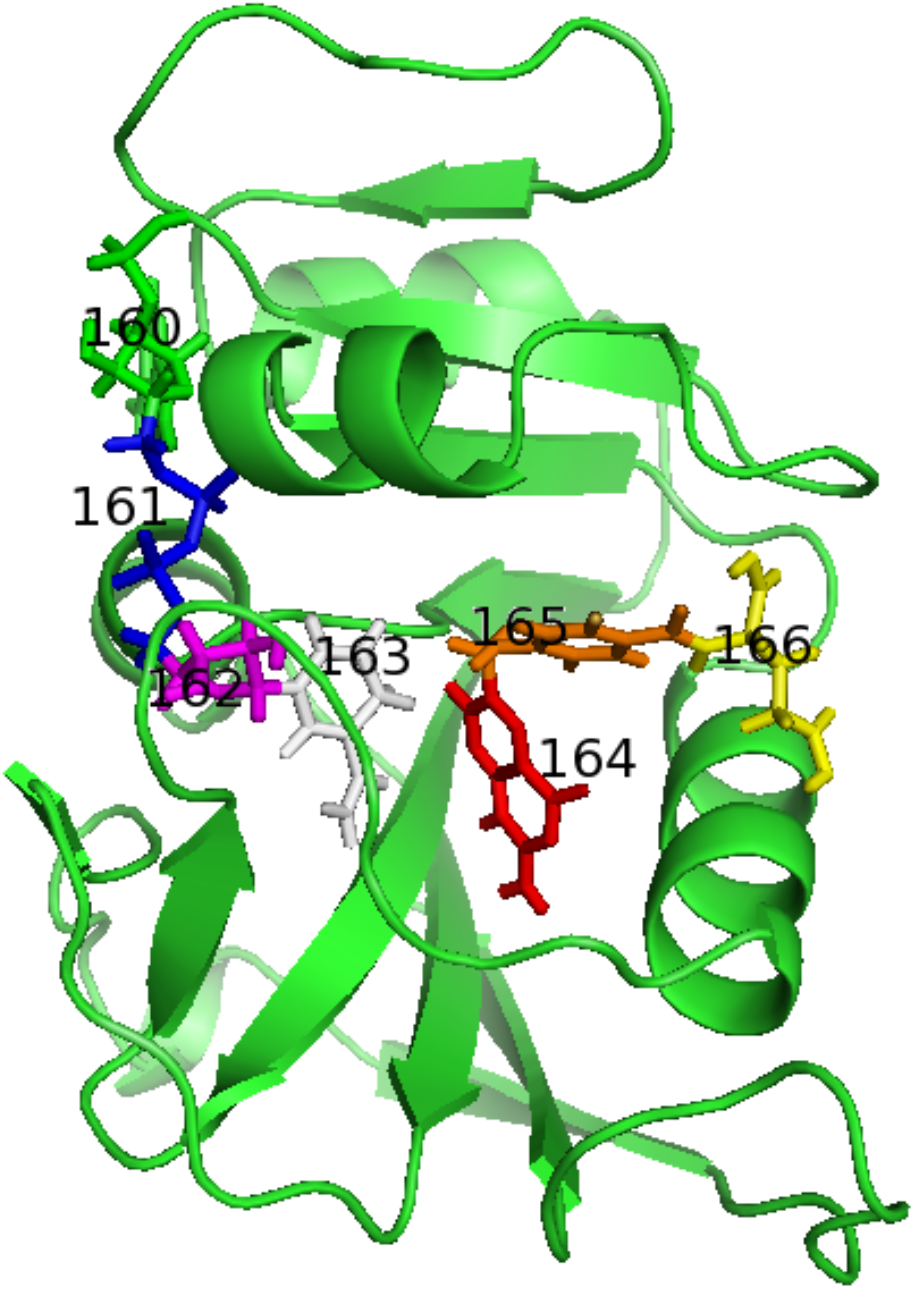
The coarse-grained NADPH and dihydrofolate. The NADPH is divided into four groups, 160-163, colored by green, blue, purple and white, respectively. The dihydrofolate is divided into three groups 164, 165 and 166 colored by red, orange and yellow.

**Figure S3.**
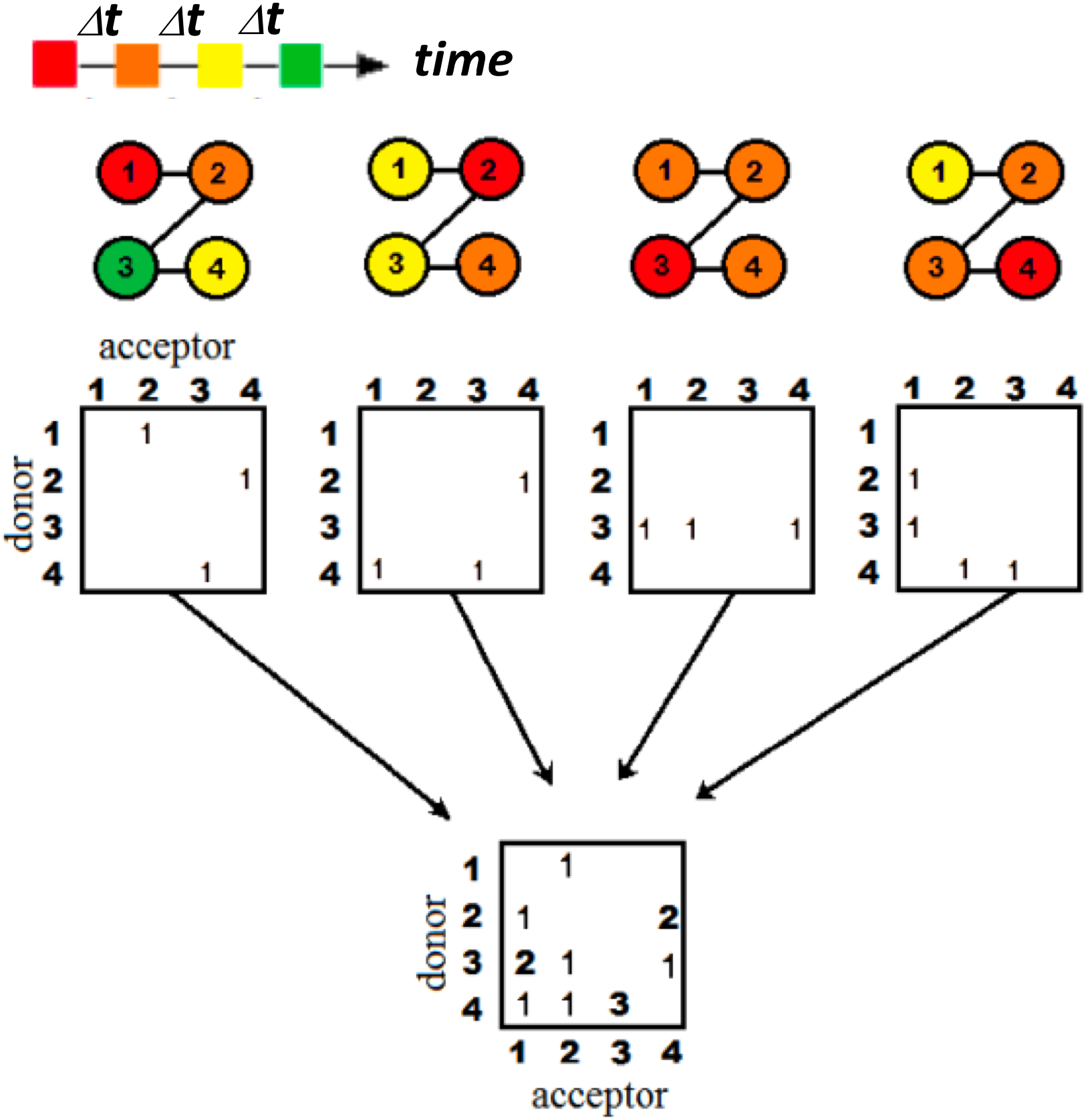
Recording the signals into the connection matrices. The colors indicate a time sequence of characteristic times. The circles with number linked by solid line are the model protein sequences, where the number is the atomic index. Given different perturbed forces,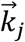, the signals propagation pathways were varied. The squares are the matrices used to record the signals. A connected pair satisfying criteria was recorded into an element of a connection matrix by adding a count. Following eq. (2), after summation over all matrices with different 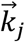, we have the full-atom connected matrix *F*(*a*_*d*_, *a*_*r*_).

**Figure S4.**
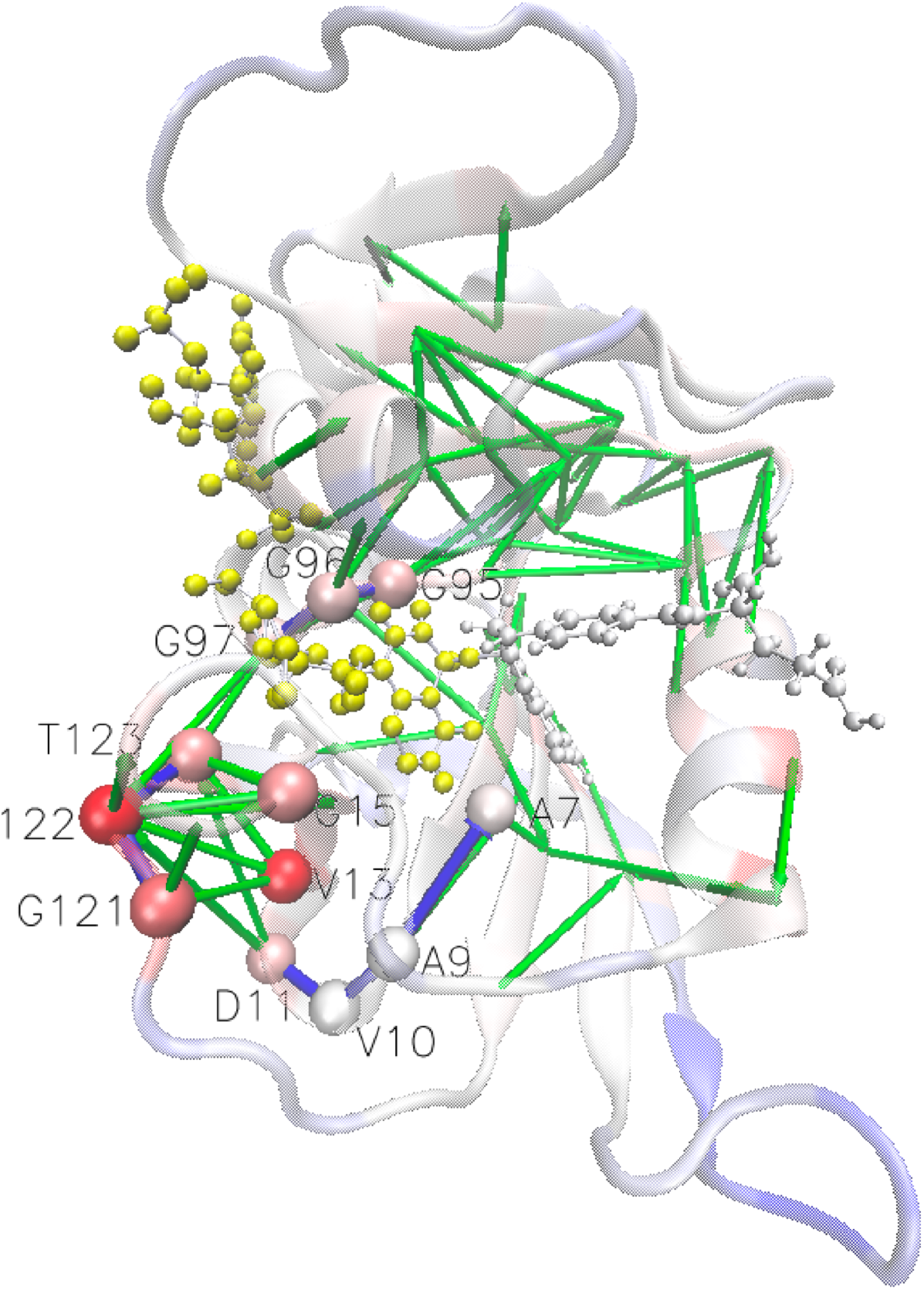
The communication pathways from a remote allosteric site G121 to sites close to catalytic centers. The pathway through the upper-middle (in blue) G121(0.88) →D122(1.00) →T123(0.81) → G97(0.70) → G96(0.73) →G95(0.76) ; The pathway through underneath the cofactor (in blue) : G121(0.88) → D122(1.00) →D11(0.75) → V10 (0.60) →A9(0.63) →A7(0.67). The values in the parentheses are the communication scores (CSs); the direction of signal propagation from one residue (donor) to the other (acceptor) in a residue pair can be assigned is because the communication is asymmetric in CGCM, where *C*(*R*_*i*_, *R*_*j*_) *≠ C*(*R*_*j*_, *R*_*i*_) (although the two numbers are usually close). The direction of signal is said from residue *i* to *j* if *C*(*R*_*i*_, *R*_*j*_) *> C*(*R*_*j*_, *R*_*i*_).

**Figure S5.**
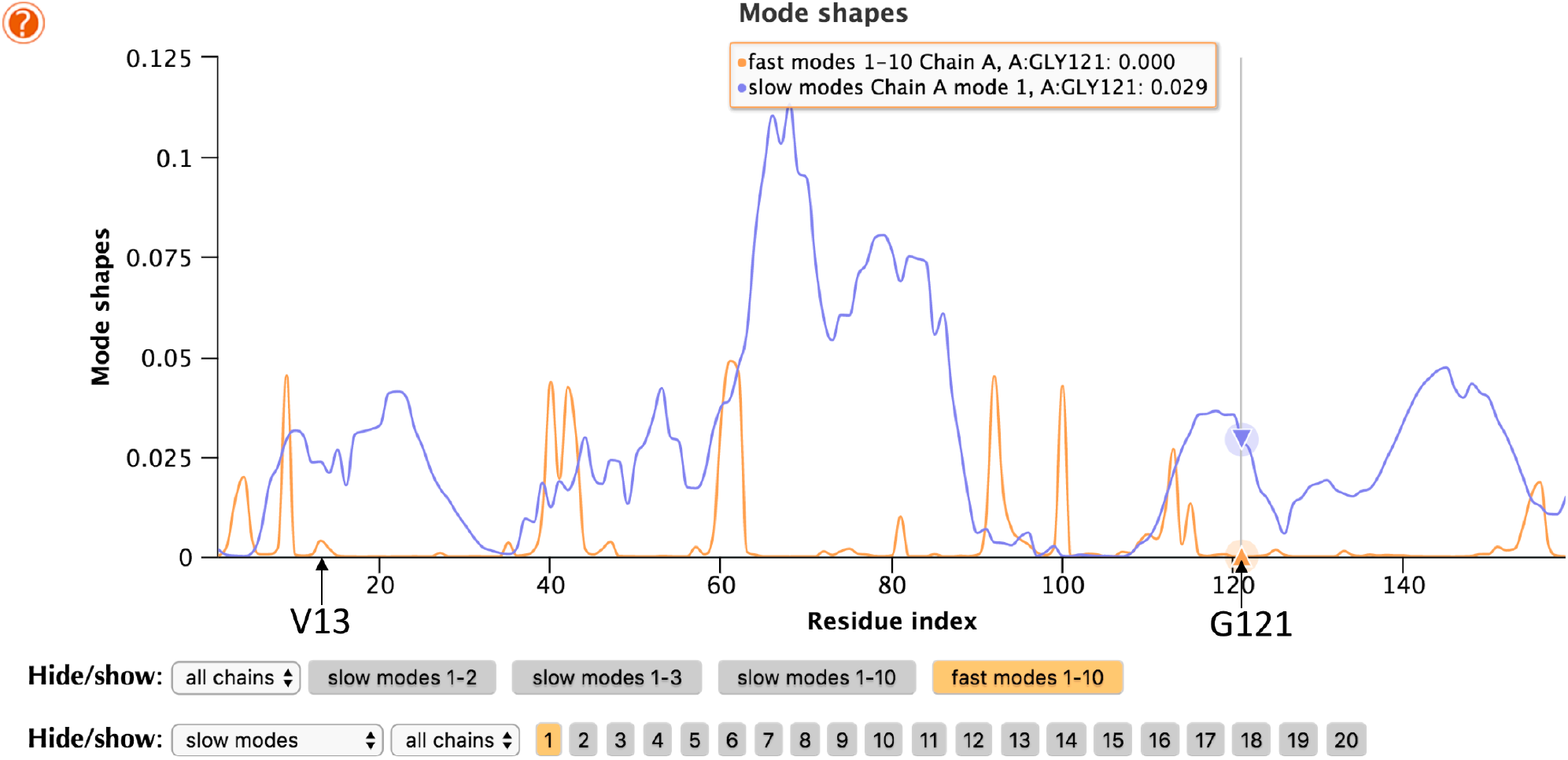
The ENM slowest mode (orange) and the average of the fastest 10 ENM mode (blue) for ternary DHFR complex (1RX6) are plotted against residue index. Frequent communicators V13-G121 are marked by black arrows. The analysis and resulting figure are taken from the DynOmics website (Li et al., 2017; https://dyn.life.nthu.edu.tw/oENM/).

**Table S1.** The communication scores (CSs) of residues in the DHFR

| rid / score |  | rid / score |  | rid / score |  | rid / score |  | rid / score |  | rid / score |  | rid / score |  | rid / score |  |
| --- | --- | --- | --- | --- | --- | --- | --- | --- | --- | --- | --- | --- | --- | --- | --- |
| **1** | 0.72 | **21** | 0.73 | **41** | 1.14 | **61** | 1.18 | **81** | 0.88 | **101** | 0.96 | **121** | 1.41 | **141** | 0.39 |
| **2** | 0.76 | **22** | 0.94 | **42** | 1.07 | **62** | 1.03 | **82** | 0.92 | **102** | 0.88 | **122** | 1.52 | **142** | 0.33 |
| **3** | 1.04 | **23** | 0.81 | **43** | 1.02 | **63** | 0.96 | **83** | 0.84 | **103** | 0.85 | **123** | 1.29 | **143** | 0.49 |
| **4** | 1.22 | **24** | 0.98 | **44** | 1 | **64** | 0.88 | **84** | 0.81 | **104** | 0.79 | **124** | 0.86 | **144** | 0.33 |
| **5** | 1.03 | **25** | 1.06 | **45** | 1.12 | **65** | 0.66 | **85** | 0.83 | **105** | 0.88 | **125** | 1.08 | **145** | 0.58 |
| **6** | 1.1 | **26** | 1.31 | **46** | 1.14 | **66** | 0.59 | **86** | 0.76 | **106** | 0.92 | **126** | 0.93 | **146** | 0.54 |
| **7** | 1.08 | **27** | 0.94 | **47** | 1.06 | **67** | 0.69 | **87** | 0.66 | **107** | 1.02 | **127** | 0.81 | **147** | 0.74 |
| **8** | 1.03 | **28** | 0.83 | **48** | 1.12 | **68** | 0.68 | **88** | 0.72 | **108** | 0.7 | **128** | 0.64 | **148** | 0.74 |
| **9** | 0.98 | **29** | 1.31 | **49** | 0.66 | **69** | 0.72 | **89** | 0.59 | **109** | 0.97 | **129** | 0.67 | **149** | 0.7 |
| **10** | 0.93 | **30** | 0.85 | **50** | 0.73 | **70** | 0.69 | **90** | 0.94 | **110** | 1.22 | **130** | 0.54 | **150** | 0.87 |
| **11** | 1.12 | **31** | 1.22 | **51** | 0.85 | **71** | 0.69 | **91** | 1.01 | **111** | 1.23 | **131** | 0.63 | **151** | 0.89 |
| **12** | 0.91 | **32** | 1 | **52** | 0.61 | **72** | 0.83 | **92** | 1.17 | **112** | 0.94 | **132** | 0.63 | **152** | 0.98 |
| **13** | 1.52 | **33** | 0.78 | **53** | 1.06 | **73** | 1.07 | **93** | 1.11 | **113** | 1.08 | **133** | 0.7 | **153** | 0.94 |
| **14** | 1.22 | **34** | 1.12 | **54** | 1.01 | **74** | 0.87 | **94** | 1.18 | **114** | 0.98 | **134** | 0.72 | **154** | 0.85 |
| **15** | 1.29 | **35** | 1.21 | **55** | 0.81 | **75** | 0.95 | **95** | 1.21 | **115** | 0.98 | **135** | 0.84 | **155** | 1.23 |
| **16** | 1.01 | **36** | 0.79 | **56** | 1.03 | **76** | 0.7 | **96** | 1.14 | **116** | 0.9 | **136** | 0.61 | **156** | 0.9 |
| **17** | 1.17 | **37** | 1.05 | **57** | 1.03 | **77** | 0.84 | **97** | 1.07 | **117** | 0.9 | **137** | 0.68 | **157** | 0.85 |
| **18** | 0.92 | **38** | 1.12 | **58** | 1.08 | **78** | 0.82 | **98** | 0.97 | **118** | 0.71 | **138** | 1.02 | **158** | 0.64 |
| **19** | 0.97 | **39** | 1.18 | **59** | 0.89 | **79** | 0.92 | **99** | 1.02 | **119** | 0.78 | **139** | 0.71 | **159** | 0.72 |
| **20** | 0.97 | **40** | 1.21 | **60** | 1.07 | **80** | 0.95 | **100** | 1.09 | **120** | 0.67 | **140** | 0.65 |  |  |

The bold texts indicate the residue indices (rid).

**Table S2.** The ConSurf scores of residues in DHFR

| ind / score |  | ind / score |  | ind/ score |  | ind / score |  | ind / score |  | ind / score |  | ind / score |  | ind / score |  |
| --- | --- | --- | --- | --- | --- | --- | --- | --- | --- | --- | --- | --- | --- | --- | --- |
| **1** | 7 | **21** | 8 | **41** | 6 | **61** | 8 | **81** | 8 | **101** | 1 | **121** | 8 | **141** | 7 |
| **2** | 7 | **22** | 9 | **42** | 9 | **62** | 4 | **82** | 3 | **102** | 5 | **122** | 9 | **142** | 1 |
| **3** | 8 | **23** | 6 | **43** | 9 | **63** | 8 | **83** | 1 | **103** | 3 | **123** | 8 | **143** | 5 |
| **4** | 4 | **24** | 9 | **44** | 9 | **64** | 8 | **84** | 1 | **104** | 6 | **124** | 1 | **144** | 8 |
| **5** | 8 | **25** | 9 | **45** | 8 | **65** | 3 | **85** | 4 | **105** | 6 | **125** | 9 | **145** | 5 |
| **6** | 7 | **26** | 6 | **46** | 9 | **66** | 1 | **86** | 1 | **106** | 1 | **126** | 9 | **146** | 3 |
| **7** | 9 | **27** | 9 | **47** | 1 | **67** | 1 | **87** | 1 | **107** | 8 | **127** | 3 | **147** | 8 |
| **8** | 6 | **28** | 8 | **48** | 7 | **68** | 1 | **88** | 1 | **108** | 5 | **128** | 1 | **148** | 1 |
| **9** | 8 | **29** | 5 | **49** | 9 | **69** | 4 | **89** | 1 | **109** | 5 | **129** | 2 | **149** | 5 |
| **10** | 1 | **30** | 6 | **50** | 8 | **70** | 1 | **90** | 8 | **110** | 7 | **130** | 1 | **150** | 1 |
| **11** | 8 | **31** | 9 | **51** | 9 | **71** | 6 | **91** | 4 | **111** | 7 | **131** | 1 | **151** | 5 |
| **12** | 5 | **32** | 9 | **52** | 7 | **72** | 5 | **92** | 7 | **112** | 6 | **132** | 1 | **152** | 3 |
| **13** | 7 | **33** | 3 | **53** | 9 | **73** | 1 | **93** | 6 | **113** | 9 | **133** | 8 | **153** | 8 |
| **14** | 9 | **34** | 2 | **54** | 9 | **74** | 6 | **94** | 8 | **114** | 1 | **134** | 1 | **154** | 1 |
| **15** | 9 | **35** | 9 | **55** | 9 | **75** | 6 | **95** | 9 | **115** | 8 | **135** | 3 | **155** | 5 |
| **16** | 4 | **36** | 5 | **56** | 6 | **76** | 3 | **96** | 9 | **116** | 4 | **136** | 3 | **156** | 5 |
| **17** | 5 | **37** | 8 | **57** | 9 | **77** | 8 | **97** | 8 | **117** | 4 | **137** | 5 | **157** | 2 |
| **18** | 9 | **38** | 7 | **58** | 3 | **78** | 2 | **98** | 4 | **118** | 1 | **138** | 2 | **158** | 9 |
| **19** | 1 | **39** | 7 | **59** | 9 | **79** | 1 | **99** | 6 | **119** | 5 | **139** | 7 | **159** | 7 |
| **20** | 8 | **40** | 7 | **60** | 8 | **80** | 1 | **100** | 8 | **120** | 3 | **140** | 1 |  |  |

The bold texts indicate the residue indices (ind).

**Table S3.** Contribution per perturbation site to C(13,121) and C(13,122)

| **perturbed sites** | **C(13,121)** | **C(13,122)** |
| --- | --- | --- |
| **5** | 0.0788 | 0.1015 |
| **6** | 0.0606 | 0.0415 |
| **7** | 0.0673 | 0.0757 |
| **14** | 0.0428 | 0.0265 |
| **20** | 0.0046 | 0.0114 |
| **22** | 0.0271 | 0.0134 |
| **27** | 0.0646 | 0.0836 |
| **28** | 0.1159 | 0.1455 |
| **31** | 0.0508 | 0.0252 |
| **46** | 0.0904 | 0.0837 |
| **49** | 0.0195 | 0.039 |
| **50** | 0.012 | 0.0349 |
| **54** | 0.0816 | 0.0905 |
| **94** | 0.0458 | 0.1476 |
| **95** | 0.0713 | 0.0745 |
| **96** | 0.0315 | 0.0401 |
| **100** | 0.1662 | 0.1053 |
| **162** | 0.1086 | 0.0932 |
| **163** | 0.1028 | 0.1435 |
| **164** | 0.1687 | 0.1328 |
| **165** | 0.0073 | 0.0107 |
| **Total score** | 1.4184 | 1.5201 |

## SUPPORTING METHODS

### Time-independent linear response theory

As described in our previous work (Yang et al., 2014; Ref 49), the time trajectory of changes in atomic position *i* can be described by the interplay between a velocity-position time correlation function and time-dependent perturbation forces summed over the atoms that are coupled with the atom *i*(54).

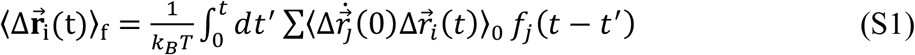

where 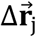 is the deviation from the mean of atom *j* due to the external force 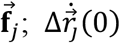 is the velocity of atom *j*; *k*_*B*_ and *T* are the Boltzmann constant and temperature, respectively, 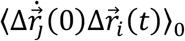 is the velocity-position time-correlation function sampled in the absence of perturbations (noted by subscript “0*”*), which can be expressed in the normal-mode space, where modes are treated as independent 1-D harmonic oscillators under solvent damping using the Langevin equation^2^. The detail derivation is referred to our previous work(54)(54b).

## The time-dependent linear response theory using impulse forces (IF-tdLRT)

Substitute the time-dependent force *f*_*j*_(*t – t*^*’*^) in eq. (S1) with an impulse force (54)(54b), presented by a delta function 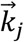, the format for the IF-tdLRT is

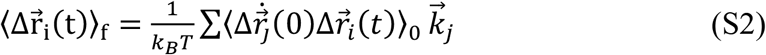

Per Chandrasekhar’s description of damped harmonic oscillators^2^, when *2ω*_*m*_ *< β*, it can be derived that

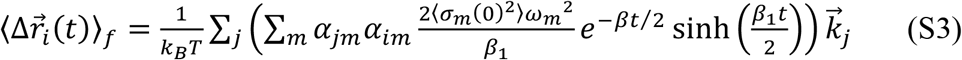

when 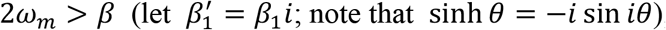, it is

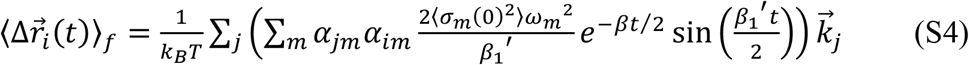

## Notes

### Competing Interest Statement

The authors have declared no competing interest.

### Summary of Updates

This is an important update where we have added new sections on MD, biochemical, cell line results of allosteric inhibitors against the tumorigenic autophagin protein ATG4B.

